# The genetic ancestry of African, Latino, and European Americans across the United States

**DOI:** 10.1101/009340

**Authors:** Katarzyna Bryc, Eric Y. Durand, J. Michael Macpherson, David Reich, Joanna L. Mountain

## Abstract

Over the past 500 years, North America has been the site of ongoing mixing of Native Americans, European settlers, and Africans brought largely by the Trans-Atlantic slave trade, shaping the early history of what became the United States. We studied the genetic ancestry of 5,269 self-described African Americans, 8,663 Latinos, and 148,789 European Americans who are 23andMe customers and show that the legacy of these historical interactions is visible in the genetic ancestry of present-day Americans. We document pervasive mixed ancestry and asymmetrical male and female ancestry contributions in all groups studied. We show that regional ancestry differences reflect historical events, such as early Spanish colonization, waves of immigration from many regions of Europe, and forced relocation of Native Americans within the US. This study sheds light on the fine-scale differences in ancestry within and across the United States, and informs our understanding of the relationship between racial and ethnic identities and genetic ancestry.

## Introduction

Over the last several hundred years, the United States has been the site of ongoing mixing of peoples of continental populations that were previously separated by geography. Native Americans, European immigrants to the Americas, and Africans brought to the New World largely via the Trans-Atlantic slave trade were brought together in the New World. Mating between individuals with different continental origins, which we refer to here as “population admixture”, results in individuals who carry DNA inherited from multiple populations. While US government census surveys and other studies of households in the US have established fine-scale self-described ethnicity at the state and county level^1^, the relationship between genetic ancestry and self-reported ancestry for each region has not been deeply characterized. Understanding genetic ancestry of individuals from a self-reported population, and differences in ancestry patterns among regions, can inform medical studies and personalized medical treatment^2^. The genetic ancestry of individuals can also shed light on the history of admixture and migrations within different regions of the US, which is of interest to historians and sociologists.

Previous studies have shown that African Americans in the US typically carry segments of DNA shaped by contributions from peoples of Europe, Africa, and the Americas, resulting in variation in African and European admixture proportions across individuals and differences in groups across parts of the country^3–5^. More recent studies using high-density genotype data provided reliable individual ancestry estimates, illustrated the large variability in African and European ancestry proportions at an individual level, and were able to detect low proportions of Native American ancestry^6, 7, 5, 4, 8–12^. Latinos across the Americas have differing proportions of Native American, African, and European genetic ancestry, shaped by local historical interactions with migrants brought by the slave trade, European settlement, and indigenous Native American populations^13–19^, and individuals from countries across South America, the Caribbean, and Mexico have different profiles of genetic ancestry molded by each population’s unique history and interactions with local Native American populations^20–26^, ^25, 26,^ ^2^. European Americans are often used as proxies for Europeans in genetic studies ^27^. European Americans, however, have a history of admixture of many genetically distinct European populations^28, 29^. Studies have shown evidence that European Americans also have non-European ancestry, including African, Native American and Asian ancestry, though it has been poorly quantified with some discordance among estimates even within studies^30–33^.

That genetic ancestry of self-described groups varies across geographic locations in the US has been documented in anecdotal examples, but has not previously been explored systematically. Most early studies of African Americans have limited resolution of ancestry due to small sample sizes and few genetic markers, while recent studies typically have limited geographic scope. Though much work has been done to characterize the genetic diversity among Latino populations from across the Americas, it is unclear the extent to which Latinos within the US share or mirror these patterns on a national or local scale. Most analyses have relied on mitochondrial DNA, Y chromosomes, or small sets of ancestry informative markers, and few high-density genome-wide SNP studies have explored fine-scale patterns of African and Native American ancestry in individuals living across the US.

We perform the first large-scale, nationwide study of African Americans, Latinos, and European Americans using high-density genotype data to examine suble ancestry patterns in these three groups across the US. To improve the understanding of the relationship between genetic ancestry and self-reported ethnic and racial identity, and to characterize heterogeneity in the fine-scale genetic ancestry of groups from different parts of the US, we inferred the genetic ancestry of 5, 269 self-reported African Americans, 8, 663 Latinos, and 148, 789 European Americans, who are 23andMe customers living across the US, using high-density SNPs genotype data from 650K to 1M arrays. 23andMe customers take an active role in participating in research by submitting saliva samples, consenting for data to be used for research, and completing surveys. We generated cohorts of self-reported European American, African American, and Latino individuals from self-reported ethnicity and identity. We obtain ancestry estimates from genotype data using a Support Vector Machine-based algorithm that infers population ancestry using reference panels to investigate patterns of Native American, African, and European ancestry in all three groups, leveraging geographic information collected through surveys (Durand *et al.*, in prep). For details on genotyping and *Ancestry Composition* methods, see *Methods*.

## Results and Discussion

### Patterns of genetic ancestry of self-reported African Americans

We find systematic differences between states in mean ancestry proportions of self-reported African Americans. In particular, we find differences between states that were slave-holding up to the time of the Civil War and “free” states in the US. Consistent with previous studies^3,^ ^34^, the diversity of ancestry profiles of 23andMe African Americans reveal that individuals comprise the full range from 0% to 100% African ancestry, but, further, that there are differences in estimates of ancestry proportions among regions. On average the highest levels of African ancestry are found in African Americans from the South, especially South Carolina and Georgia (Figure 1A). We find lower proportions of African ancestry in the Northeast, the Midwest, the Pacific Northwest and California. For mean ancestry proportions and sample sizes for states and regions, refer to Supplementary Tables S2 and S3. The small sample sizes from some areas of the US, including parts of the Midwest and Mountain regions reflects the lower population density of African Americans residing in these regions^35^. Reflected in these ancestry patterns are migration routes, such as the route through important Southern seaports bringing Africans through the trans-Atlantic slave trade^36, 37^.

**Figure 1:**
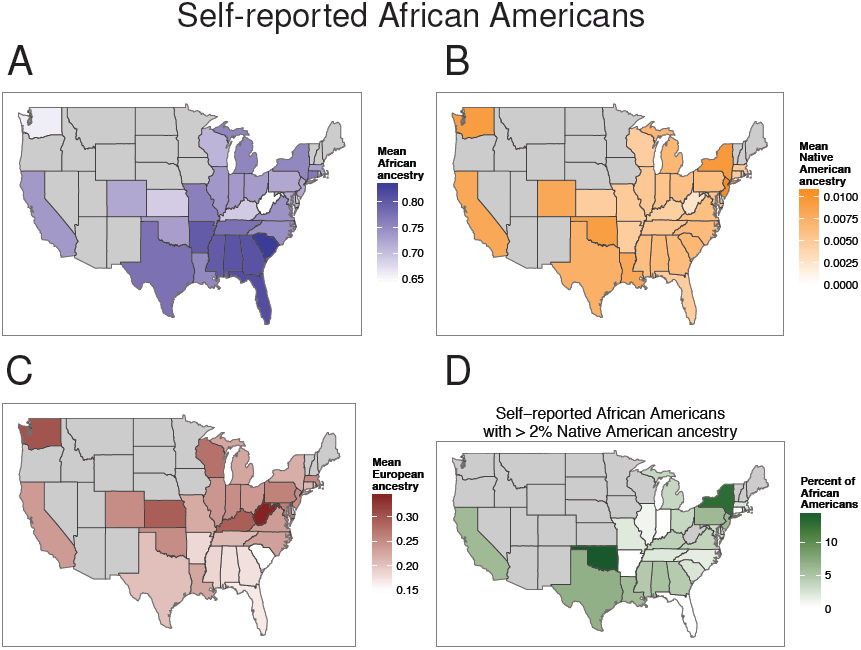
The distribution of ancestry of self-reported African Americans across the US. (A) Differences in levels of African ancestry in African Americans (blue) (B) Differences in levels of Native American ancestry in African Americans (orange) (C) Differences in levels of European ancestry of African Americans (red), from each state. States with fewer than 10 individuals are excluded in gray. (D) The geographic distribution of self-reported African Americans with Native American ancestry. The proportion of African Americans in each state who have 2% or more Native American ancestry is shown by shade of green. States with fewer than 20 individuals are excluded in gray.

Genome-wide ancestry estimates of African Americans show average proportions of 73.2% African, 24.0% European, and 0.75% Native American ancestry. Though mean estimates of Native American ancestry are low, many African Americans carry detectable levels of Native American ancestry. Over 5% of African Americans are estimated to carry at least 2% Native American ancestry genome-wide (Figures S1 and 1D). Consistent with historical narratives and family histories, our estimate suggests that 1 in every 20 African Americans carries Native American ancestry, a higher rate than we detected in self-reported European Americans. An individual that carries more than 2% Native American ancestry can arise from one genetically Native American ancestor within the last 9 or so generations, or multiple genealogical Native American ancestors^38^. Using a lower threshold of 1% Native American ancestry, we estimate that about 22% of African Americans carry some Native American ancestry, which implies that more than 1 in 5 African Americans may have a Native American ancestor in their genealogy within the last 11 generations (Figure S2).

Even excluding individuals with no African ancestry, which are likely the result of survey errors, we still estimate a higher European, and corresponding lower African, mean genetic ancestry proportion in 23andMe African Americans compared to previous studies of African Americans. A significant difference between the 23andMe cohort of African Americans and many groups previously studied is geographic sampling location. Our cohort reflects heavier sampling from California and New York, likely driven by population density as well as awareness of genetic testing or 23andMe. Both are regions where African Americans have lower mean African ancestry than other studies of African Americans, which are often drawn from locations in the South. However, participation in 23andMe is not free and requires online access, therefore it is important to note that other social, cultural, or economic factors may interact to affect ancestry proportions of those individuals who choose to participate in 23andMe.

The amount of Native American ancestry estimated for African Americans varies across states in the US. African Americans in the West and Southwest on average carry higher levels of Native American ancestry, a trend that is largely driven by individuals with less than 2% Native American ancestry (Figure 1B) rather than by many individuals with substantial ancestry (Figure S2). Oklahoma shows the highest proportion of African Americans with substantial Native American ancestry, where over 14% of African Americans from Oklahoma carry at least 2% Native American ancestry (Figures 1B and S2). Oklahoma was the site of contact between Native Americans and African Americans following the Trail of Tears migration in the 1830’s^39, 40^, where black slaves comprised a significant part of the population in the 1860’s^41^, and the location of the slave-holding “Five Civilized Tribes”. In contrast, we do not observe higher rates of Native American ancestry in African Americans in Florida, which is potentially notable in light of the known history of Seminole intermarriage with blacks according to the 1860 U.S. Census^41^.

A sex bias in African American ancestry, with greater male European and female African contributions, has been suggested through mtDNA, Y chromosome, and autosomal studies^7^. Through comparison of estimates of X chromosome and genome-wide African and European ancestry proportions, we estimate that approximately 6% of ancestors of African Americans were European females, while 19% were European males. On average across African Americans, we estimate that the X chromosome has a 5% increase in African ancestry and 18% reduction in European ancestry relative to genome-wide estimates (see Table 1). Sex bias in ancestry contributions may have been driven by unbalanced sex ratios in immigration frontier settings^42^, exploitation^43^, or other social factors.

**Table 1:**
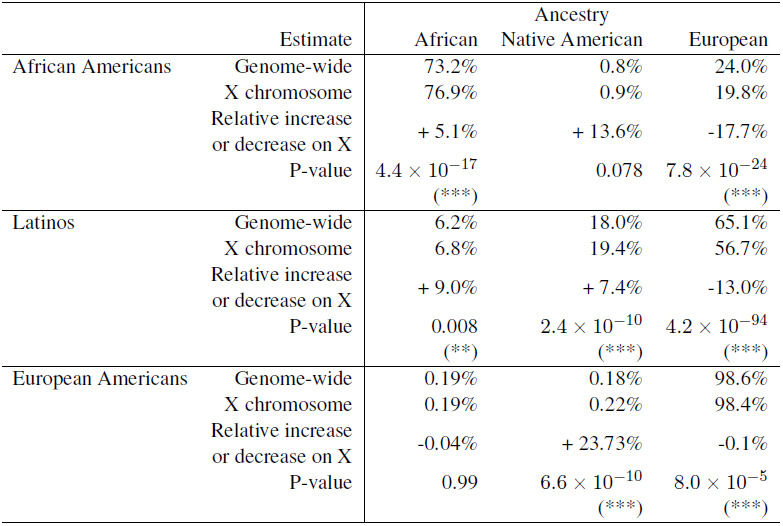
Comparison of genome-wide ancestry estimates and X chromosome estimates in African Americans, Latinos, and European Americans. Mean estimates of African, Native American, and European ancestry are shown. P-values provided are calculated using two-sided students’ t-test on individual ancestry estimates for each cohort per ancestry, with no multiple testing correction. Significance is assigned as p-value < 0.05 (*), p-value < 0.01 (**) and p-value < 0.001 (***). Relative increase on the X chromosome is calculated as the absolute difference, X chromosome estimate minus genome-wide estimate, divided by the genome-wide estimate.

|  |  | Ancestry |  |  |
| --- | --- | --- | --- | --- |
|  | Estimate | African | Native American | European |
| African Americans | Genome-wide | 73.2% | 0.8% | 24.0% |
|  | X chromosome | 76.9% | 0.9% | 19.8% |
|  | Relative increase or decrease on X | + 5.1% | + 13.6% | -17.7% |
| | P-value | $4.4 \times 10^{-17}$<br>(***) | 0.078 | $7.8 \times 10^{-24}$<br>(***) |
| Latinos | Genome-wide | 6.2% | 18.0% | 65.1% |
|  | X chromosome | 6.8% | 19.4% | 56.7% |
|  | Relative increase or decrease on X | + 9.0% | + 7.4% | -13.0% |
| | P-value | 0.008<br>(**) | $2.4 \times 10^{-10}$<br>(***) | $4.2 \times 10^{-94}$<br>(***) |
| European Americans | Genome-wide | 0.19% | 0.18% | 98.6% |
|  | X chromosome | 0.19% | 0.22% | 98.4% |
|  | Relative increase or decrease on X | -0.04% | + 23.73% | -0.1% |
| | P-value | 0.99 | $6.6 \times 10^{-10}$<br>(***) | $8.0 \times 10^{-5}$<br>(***) |

We used the lengths of segments of European, African, and Native American ancestry to estimate a best-fit model of admixture history among these populations for African Americans (Figure S3) using the demographic modeling of *TRACTS*^44^. We fit a two-pulse model of admixture among three ancestral populations to estimate Native American admixture 12 generations ago in the past, with more recent African/European admixture 3.5 or 4 generations ago. These dates are estimated as the best fit for a pulse admixture event: since they represent an average over more continuous or multiple migrations, initial admixture is likely to have begun earlier.

We find that our estimates of sex bias in ancestry contributions in African Americans, with greater male European ancestry and greater female African contributions, support over three times as many European male ancestors as female European ancestors in African Americans. Sex bias in ancestry contributions has been suggested to stem from interactions with male European slave-owners and female African slaves^45^. However, our estimates of admixture dates, with a mean admixture date of 4 generations, implies that the majority of admixture events between Europeans and African ancestry has taken place much more recently than 1865, and in particular, may have taken place after slavery. Taken together, these datapoints suggest that sex biased admixture may also reflect post-slavery social and cultural influences.

### Patterns of genetic ancestry of self-reported Latinos

Latinos encompass nearly all possible combinations of African, Native American, and European ancestries, with the exception of individuals who have a mix of African and Native American ancestry without European ancestry (see Figure S4A and S1). On average, we estimate that Latinos in the US carry 18.0% Native American ancestry, 65.1% European ancestry, and 6.2% African ancestry. We find the highest levels of estimated Native American ancestry in self-reported Latinos from states in the Southwest, especially those close to Mexico (Figure 2C). We find the highest mean levels of African ancestry in Latinos highest in Louisiana, some states in the South, and states in the Midwest and Atlantic (Figure 2A).

**Figure 2:**
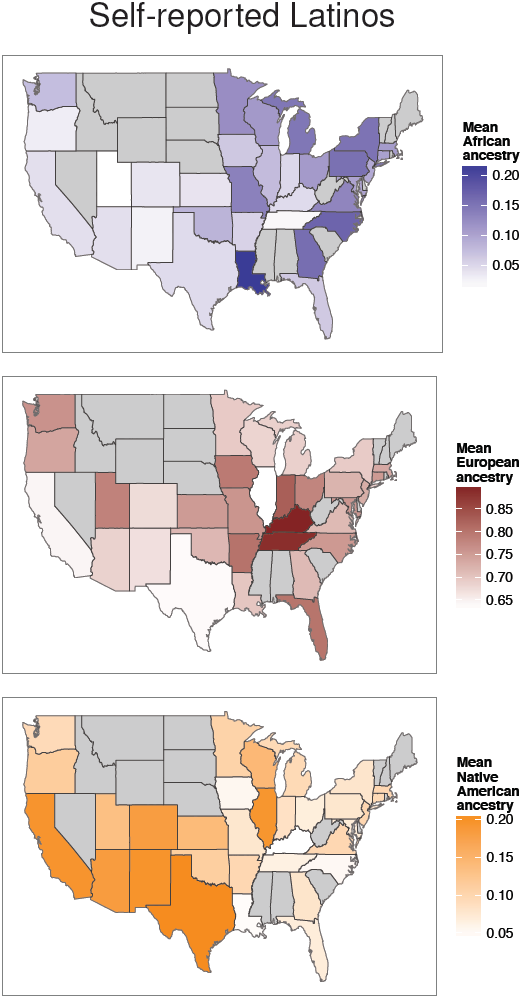
The distribution of ancestry of self-reported Latinos across the US. Differences in mean levels of African, European, and Native American ancestry in Latinos from each state is shown by shade of blue, red, and orange, respectively. States with fewer than 10 individuals are excluded in gray.

Further stratification of individuals by their self-reported population affiliation (e.g., “Mexican”, “Puerto Rican”, or “Dominican”) reveals a diversity in genetic ancestry, consistent with previous work studying these populations (see Figure S5 and Table S4)^21, 25, 46, 11, 47,^ ^26^. We find that Latinos who, besides reporting as “Hispanic”, also self-report as Mexican or Central American, carry more Native American ancestry than Latinos overall, while those who self-report as black, Puerto Rican, or Dominican have higher levels of African ancestry, and those who additionally self-report as white, Cuban, or South American have on average higher levels of European ancestry. These history-dependent ancestry differences are likely to underpin regional differences in ancestry proportions across states.

Consistent with previous studies that show a sex bias in admixture in Latino populations, with greater male European and female Native American ancestry^13–19^, we estimate 13% less European ancestry on the X chromosome than genome-wide (Table 1) showing proportionally greater European ancestry contributions from males. We infer elevated African and Native American ancestry on the X chromosome, corresponding to higher female ancestry contributions from both Africans and Native Americans. Our estimates of ancestry on the X chromosome provide strong evidence for more male European ancestors and more female non-European (both African and Native American) ancestors in Latinos living in the US.

Latinos show higher proportions of inferred Iberian ancestry than both European Americans and African Americans (Figure S6). We estimate that Iberian ancestry composes as much as a third of the European ancestry in Latinos in Florida, New Mexico, and other parts of the Southwest, likely either reflecting early Spanish influence and rule in these regions, or recent immigration from Latin America, which may also be associated with higher levels of Iberian ancestry in New York and New Jersey. Regions with higher Iberian ancestry also correspond to regions with greater Native American ancestry; disentangling whether higher levels of Native American ancestry in the Southwest reflects the legacy of indigenous Native American ancestors or is the result of recent Latino immigrants into the Southwest may be possible through future studies of admixture dating or more precise Native American ancestry delineations.

### Patterns of genetic ancestry of self-reported European Americans

We estimated proportions of British/Irish, Eastern European, Iberian, and Scandinavian ancestry (Figure 3), and other European subpopulation ancestries (Figure S7), and found strong regional differences across states reflecting known major historical migrations from Europe. Inferred British/Irish ancestry is found in European Americans from all states at mean proportions of above 20%, and represents a majority of ancestry, above 50% mean proportion, in states such as Missisippi, Arkansas, and Tennessee. We note that these states are similarly highlighted in the map of the self-reported “American” ethnicity in the US census survey^1^, which may reflect regions with lower subsequent migration from other parts of Europe. Inferred Eastern European ancestry is found at its highest levels in Illinois, Michigan, and Pennsylvania, potentially stemming from immigration during the late 19th and early 20th centuries, settling in metropolitan areas in the Northeast and Midwest. Inferred Iberian ancestry, found overall at lower mean proportions, still represents a measurable ancestry component in Florida, Louisiana, California, and Nevada, and may point to the early Spanish rule and colonization of the Americas. Scandinavian ancestry in European Americans is highly localized; most states show only trace mean proportions of Scandinavian ancestry, while it comprises a significant proportion, upwards of 10%, of ancestry in European Americans from Minnesota and the Dakotas. The distributions of the European subpopulation ancestries in European Americans illustrate that the distribution of within-European ancestry is not homogenous among individuals from different states, and instead, reflects differences in population migrations and settlement patterns within the US.

**Figure 3:**
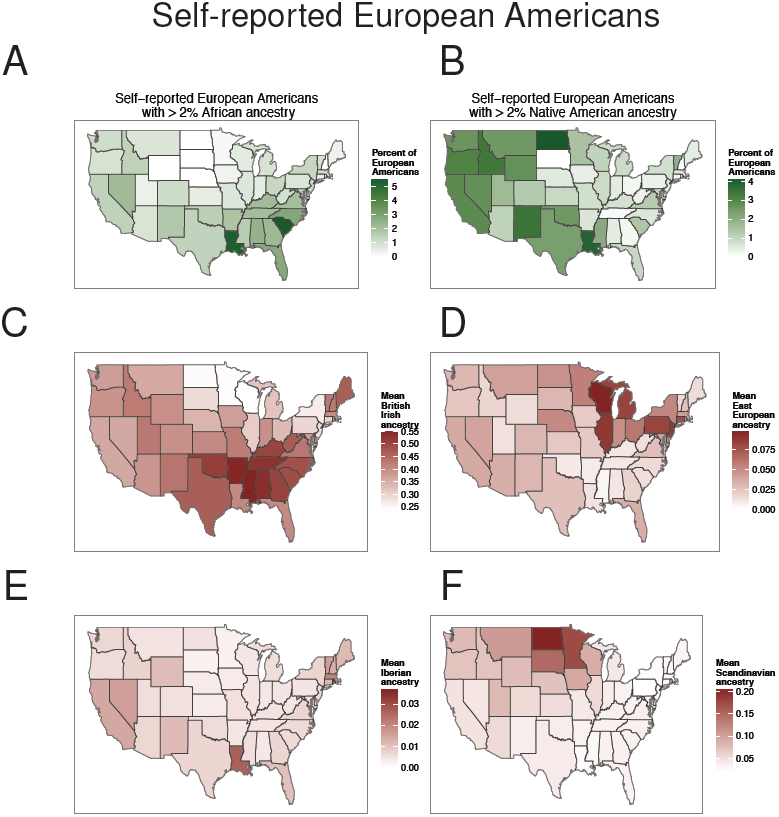
Differences in African, Native American, and European subpopulation ancestry among self-reported European Americans from different states. (A) The geographic distribution of self-reported European Americans with African ancestry. The proportion of individuals with at least 2% African ancestry, out of the total number of European Americans per state, is shown by shade of green. (B) The geographic distribution of self-reported European Americans with Native American ancestry. The proportion of European Americans who have 2% or more Native American ancestry is shown for each state. (C-F) The mean British/Irish, Eastern European, Iberian, and Scandinavian ancestry proportions among self-reported European Americans from each state are shown by shade of red.

We find that many self-reported European Americans, predominantly those living west of the Mississippi River, carry Native American ancestry (Figure 3B). Though the average levels of Native American ancestry are trace, if we examine the frequency with which European Americans carry at least 2% Native American ancestry, we see that Native American ancestry occurs most frequently in Louisiana, North Dakota, and other states in the West. We estimate that 4% of self-reported European Americans living in Louisiana and North Dakota carry segments of Native American ancestry. Using a less stringent threshold of 1%, our estimates suggest that as many as 8% of individuals from Lousiana and upwards of 3% of individuals from some states in the West and Southwest, carry Native American ancestry (Figure S8).

We estimate that a substantial fraction, at least 1.4%, of self-reported European Americans in the US carry at least 2% African ancestry. Using a less conservative threshold, approximately 3.5% of European Americans have 1% or more African ancestry (Figure S9). Consistent with previous anecdotal results^33^, the frequency of European American individuals who carry African ancestry varies strongly by state and region of the US (Figure 3A). In particular, individuals with African ancestry are found at much higher frequencies in states in the South than in other parts of the US: about 5% of self-reported European Americans living in South Carolina and Louisiana have at least 2% African ancestry. Lowering the threshold to at least 1% African ancestry (potentially arising from one African genealogical ancestor within the last 11 generations), European Americans with African ancestry comprise as much as 12% of European Americans from Louisiana and South Carolina and about 1 in 10 individuals in other parts of the South (Figure S9). Lousiana’s high levels of African ancestry in European Americans are consistent with historical accounts of intermarriage in the New Orleans area^48, 49^. Applying these estimates to self-reported ethnicity data from the 2010 US Census suggests that over 6 million Americans, who self-identify as European, may carry African ancestry. Likewise, as many as 5 million Americans who self-identify as European may have at least 1% Native American ancestry.

We find evidence that sex biased admixture processes are widespread in US history. Non-European ancestry in European Americans follows a sex bias in admixture contributions from males and females, as seen in African Americans and Latinos. The ratio between X chromosome and genome-wide Native American ancestry in European Americans shows greater Native American female and higher European male ancestry contributions. Though we do not observe evidence of a sex bias in African ancestry contributions overall, analysis of European Americans with at least one percent African ancestry reveals 15% higher African ancestry on the X relative to genome-wide estimates (p-value 0.013). This increase suggests female-African and male-European sex bias in European Americans follows the same direction as in African Americans and Latinos, with greater male European and female African and Native American contributions. Our estimates of sex bias in European Americans, as well as in African American and Latino populations, show that sex ratios on the frontier, or other social factors, have resulted in asymmetrical admixture processes in all US populations studied.

### Correlations with population density

We find that proportions of Native American and African ancestry in 23andMe customers in each state are generally significantly correlated with the population density of African Americans and Latinos in each state (Figures S10, S11, and S12). For example, levels of African ancestry in a state are highly correlated with African American state population density in European Americans and Latinos (both p-values < 10^−4^). Mean ancestry proportions of Native American ancestry in each state are correlated to Latino state population density in European Americans, and in Latinos (p-values < 10^−6^ and < 10^−2^ respectively). Given the highly significant statistics in European Americans, surprisingly, in African Americans, the correlation of African ancestry with African American population density is only marginally signficant (p-value 0.025). In African Americans, the states with the highest mean levels of African ancestry, such as South Carolina, Georgia, and Florida are not those with the highest density of African Americans. The correlation of Native American ancestry in African Americans with Latino state population density also has a marginal p-value of 0.026. Correlations between state population density and mean ancestry proportions suggest that the numbers of African and Native American individuals in a state may have shaped the ancestries of present-day individuals. Not all correlations are strongly significant, suggesting that other social or cultural factors influenced ancestry proportions, especially in African Americans.

### Relationship of self-identity and genetic ancestry

Contrary to the one-drop rule, or the “Rule of Hypodescent”, which would mandate that individuals that knowlingly carry African ancestry identify as African American, our results show that most individuals who have less than 28% African ancestry identify as European American, rather than as African American (Figures 4 and 5A). In fact, the probability of self-reporting as African American given a proportion of African ancestry follows a logistic probability curve (Figure S13, Table S5), suggesting that individuals identify roughly with the majority of their ancestry. In contrast, Latino self-reported identity becomes the most frequent identity at 5% or greater Native American ancestry (Figures S13 and 5B), suggesting differences in sociological or historical factors associated with identifying with these groups. The transitions between Latino, African American, and European American self-reported identity by proportions of African and Native American ancestry illustrate both the complexity of self-identity as well as the overlapping ancestry profiles among groups (Figure 5B).

**Figure 4:**
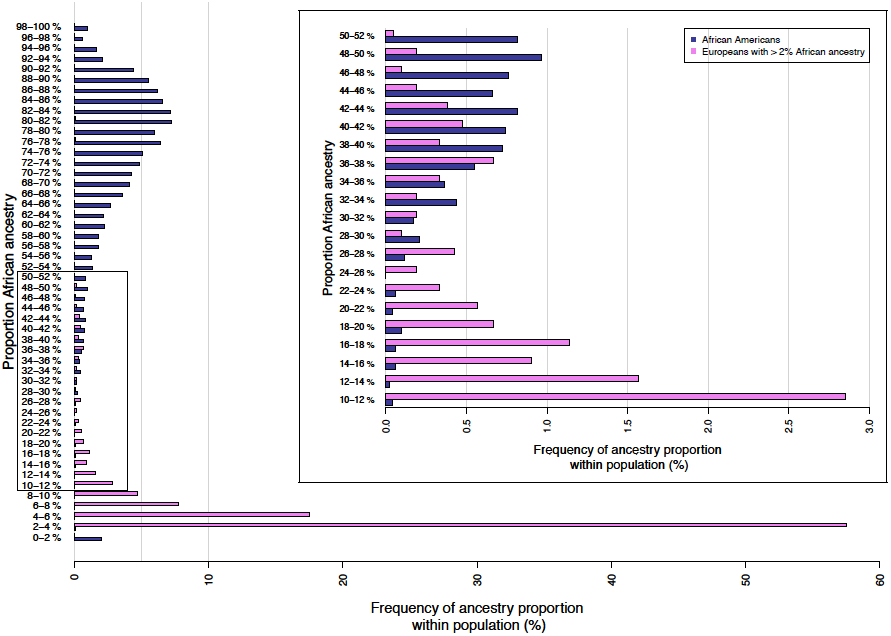
Distribution of African ancestry in African Americans and European Americans. Histogram of African Americans (blue bars), and European Americans with ≥ 2% African ancestry (violet bars). Inset: Fine-scale histogram showing the region of greatest overlap between African Americans and European Americans, where African ancestry ranges from 10% and 52%. Both histograms have been normalized for each cohort to total 100%.

**Figure 5:**
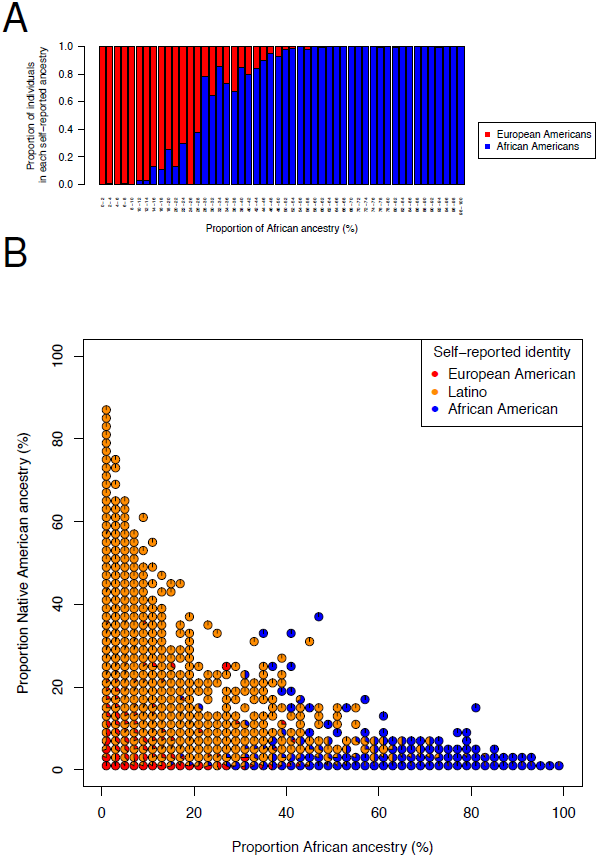
Proportions of individual self-identities by genome-wide ancestry proportions. (A) The proportion of individuals that self-report as African American versus European American for each 2% bin of African ancestry. Each vertical bar corresponds to the individuals that carry that bin of ancestry, and is colored by the proportion of African American and European American identities. Proportions are estimated from absolute numbers of individuals, not scaled by total cohort size. (B) The proportion of individuals that self-report as European American, Latino, and African American for each 2% bin of African ancestry and Native American ancestry. The proportion for each 2% bin is shown as a pie chart, with slices colored in proportion to the absolute numbers of individuals from each self-reported identity that carry those levels of genome-wide ancestry. Pie charts are omitted for bins where there were no individuals with those corresponding levels of Native American and African ancestry.

## Conclusion

### The genetic landscape of the United States

This work demonstrates that the legacy of population migrations and interactions is visible in the genetic ancestry of modern individuals living in the US. Our results suggest that genetic ancestry can be leveraged to augment historical records and inform cultural processes shaping modern populations. The relationship between self-reported identity and genetic African ancestry, as well as the low numbers of self-reported African Americans with minor levels of African ancestry, provide insight into the complexity of genetic and social consequences of racial categorization, assortative mating, and the impact of notions of ‘race’ on patterns of mating and self-identity in the US. Our results provide empirical support that, over recent centuries, many individuals with partial African and Native American ancestry have “passed” into the white community^50, 51^, with multiple lines of evidence establishing African and Native American ancestry in self-reported European Americans (see *Methods*). Though the majority of European Americans in our study did not carry Native American or African ancestry, even a small proportion of this large population that carry non-European ancestry translates into millions of European Americans who carry African and Native American ancestry. Our results suggest that the early US history, beginning in the 17th century (or around 12 generations ago), may have been a time of many population interactions resulting in admixture.

Our large numbers of individuals, high-density genotype data, and accurate and robust local ancestry estimates allow us to discern subtle differences in genetic ancestry. In spite of present-day high mobility of individuals, the genetic ancestry of present-day individuals recapitulates historical migration events, known settlement patterns, and admixture processes. Perhaps most importantly, however, our results reveal the impact of centuries of admixture in the US thereby undermining the use of cultural labels that group individuals into discrete non-overlapping bins in biomedical contexts “which cannot be adequately represented by arbitrary ‘race/color’ categories”^52^.

## Subjects and Methods

### Human Subjects

All participants were drawn from the customer base of 23andMe, Inc., a consumer personal genetics company. This dataset has been described in detail previously^53, 54^. Participants provided informed consent and participated in the research online, under a protocol approved by the external AAHRFP-accredited IRB, Ethical & Independent Review Services (E&I Review).

### Genotyping

Participants were genotyped as described previously^55^. Samples were genotyped on one of two platforms. The first is the Illumina HumanHap550+ BeadChip platform, which included SNPs from the standard HumanHap550 panel augmented with a custom set of approximately 25,000 SNPs selected by 23andMe. Two slightly different versions of this platform were used, as previously described (9). The second platform is the Illumina HumanOmniExpress+ BeadChip. This platform has a base set of 730,000 SNPs augmented with approximately 250,000 SNPs to obtain a superset of the HumanHap550+ content, as well as a custom set of about 30,000 SNPs. Customer genetic data have been previously utilized in association studies and studies of genetic relationships^56–64^.

### Research cohorts

23andMe customers were invited to fill out web-based questionnaires, including questions on ancestry and ethnicity, on state of birth, and current zip code of residence. They were also invited to allow their genetic data and survey responses to be used for research. Only data of customers who signed IRB-approved consent documents were included in our study. Survey introductions are explicit about their applications in research. For example, the ethnicity survey introduction text states that the survey responses will be used in ancestry-related research (Table S1).

#### Self-reported ancestry

It is important to note that self-reported ancestry, ethnicity, and identity (and race) are complex terms that are a result of both visible traits, such as skin color, and cultural, economic, geographical and social factors^65, 24^. As a result, the precise terminology and labels used for describing self-identity may affect survey results, and care in choice of labels should be utilized. However, we chose to maximize our available self-reported ethnicity sample size by combining information from questions asking for customer self-reported ancestry. We use survey questions to gauge responses about *identity*, which here we view as “the subjective articulation of group membership and affinity”^66^. The two questions use different nomenclature, as follows.

The first question is modeled after the US census nomenclature, and is a multi-question survey that allows for choice of “hispanic” or “not hispanic”, and participants were asked “Which of these US Census categories describe your racial identity? Please check all that apply.” from the following list of ethnicities: “white”, “black”, “american indian”, “asian”, “native hawaiian”, “other”, “not sure”, and “other racial identity”.

For inclusion into our European American cohort, individuals had to select “not hispanic” and “white”, but not any other identity. For inclusion into our Latino cohort, indiviudals had to select “hispanic”, with no other restrictions. For inclusion into our African American cohort, individuals had to select “not hispanic” and “black” and no other identity.

We also included participants that responded to a single-choice question on ethnicity, where respondents were asked to choose “What best describes your ancestry/ethnicity?” from: “african”, “african american”, “central asian”, “declined”, “east asian”, “european”, “latino”, “mideast”, “multiple ancestries”, “native american”, “not sure”, “other”, “pacific islander”, “south asian”, and “southeast asian”. Since individuals could only select one reponse, we included individuals who selected “european” in our European American cohort, those who selected “african american” in our African American cohort, and “latino” in our Latino cohort. Some African American participants included in this study were recruited through 23andMe’s Roots into the Future project www.23andme.com/roots/, aimed to increase understanding of how DNA plays a role in health and wellness, especially for diseases more common in the African American community. These individuals, who self identify as African American, Black, or African were recruited through 23andMe’s current membership, at events, and via other recruitment channels, under advisement of the Roots into the Future advisory panel (see 23andMe.com/roots/).

Lastly, when available, we excluded individuals who answered “No”, to a question whether they are living in the US. In total, our final sets included 5, 269 African Americans, 8, 663 Latinos, and 148, 789 European Americans.

### Notes on terminology and selection of populations

Throughout the manuscript, the term “Native American ancestry” refers to estimates of genetic ancestry from indigenous Americans found across North, Central, and South America, and we distinguish this term from present-day Native Americans living in the US. We use the term “Native American” to refer to indigenous peoples of the Americas, acknowledging that some people may prefer other terms such as “American Indian”.

Our estimates of African ancestry specifically aims to infer ancestry of Sub-Saharan Africa, and does not include ancestry from North Africa or the Middle East. We note that the term “Latino” has many meanings in different contexts, and in our case, we use it to refer to individuals living in the US that self-report as either “Latino” or “Hispanic”. In the present work, we do not include individuals who self-report as having multiple identities, as this represents only a small fraction of individuals in our dataset. Low rates of reporting as multi-racial or multi-ethnic is in line with previous studies; an analysis of the 2000 US Census shows that 95 percent of blacks and 97 percent of whites only acknowledge a single identity^66^. Future studies including multi-racial individuals may further illuminate patterns of genetic ancestry and the complex relationship with self-identity. Differences among states, where different levels of people self-report as mixed race may explain some regional differences in genetic ancestry. However, we note that, first, proportionally fewer people identify as mixed race than as a single identity, and second, it remains important to establish regional differences in genetic ancestry of self-reported groups even if these differences are driven, to some degree, by regional changes in self-reported identity. More work is needed to what extent regional differences are a result of how people today report their ancestry.

We note that the ancestries of 23andMe customers, and therefore the demographics of the database used for this study, largely reflect the demographics of the US, as tallied in the 2010 US census. Our study considers three cohorts that comprise the three largest self-identified groups in the US, which hence are likewise well represented in the 23andMe database. We focus on studying the distribution of European, African, and Native American ancestries and European subpopulation ancestries. These populations were selected as both being accurate to resolve and because they reflect the major waves of migration into the US just after the era of transcontinental travel began, and are found at mean frequencies of above 1% in these populations.

Our study focuses on groups and ancestries for which we are able to collect sufficient sample sizes and reference population sizes. However, we emphasize that these groups and ancestries are only a fraction of the diversity found within individuals living in the US, and as dataset sizes grow, future work should extend to include analyses of other world-wide ancestries and populations and ancestries, and their distributions across the US. Our work represents a snapshot in time of genetic ancestry and identity, and future work is needed to inform the dynamic changes and forces that shape social interactions in the past and in the future.

We note that our cohorts are likely to have ancestry from many African populations, but due to current reference sample availability, our resolution of within West African ancestries is outside the scope of our study. Likewise, our estimates of Native American ancestry arise from a summary over many distinct sub-populations, but we are limited in scope due to insufficient sample sizes from subpopulations, as we currently use individuals from Central and South American together as a reference set (see Durand *et al.* for a list of populations and sample sizes). We regret that we are, at present, unable to delve deeper into the complexity of, and sub-ancestries within, these continental populations, and do not wish to convey that within-European ancestry is of greater importance. Instead, our resolution reflects the current availability of reference individuals from different regions.

#### Validation of self-reported identity survey results

The use of self-reported data in a medical context has shown good agreement, especially in diseases that are well-defined and easily diagnosed^67–74^. 23andMe conducts internal research to validate survey data collection strategies, and traits such as self-reported age, eye color, hair color, freckles and handedness have been shown to have concordance for more than 99.25% of individuals^56^. To verify that our self-reported ethnicities were reliable, we examined the consistency of ethnicity survey responses when individuals completed both ancestry and ethnicity surveys. Since the structure of the two surveys is different, and multiple selections were allowed in one survey but not the other, we examined the replication rate of the primary ethnicity from the single-choice ethnicity survey in the multiple-selection survey.

The survey consistency was remarkably high. Out of 35, 524 self-reported “European” individuals, 35, 279 selected “white” on the ethnicity survey, yielding a per-survey error estimate of 0.2%. Out of 1, 560 self-reported “Latino” individuals, 1, 540 selected “Hispanic”, giving a per-survey error estimate of 0.7%. Lastly, out of 1, 327 self-reported “African American” individuals, 1, 287 selected “black”, resulting in a per-survey error rate estimate of 1.1%.

In addition to structural differences, the survey content used very different nomenclature, and therefore we believe our estimated error rates to be overestimates of the true error rate, since it is likely that some individuals may choose to identify with one label but not the other (ie, “African American” but not “black”). Discrepancies in the question nomenclatures is likely to increase the error rate. Furthermore, since the two surveys could be completed at different times, either before, or after, obtaining personal ancestry composition results, it is possible that viewing genetic ancestry results may have led to a change in self-reported ancestry. Such a change would be tallied as an error in our estimates, but instead reflects a true change in perceived self-identity over time. Overall, we expect that our survey data represents highly reliable ancestry information, with errors affecting fewer than 1% of of survey responses.

#### Geographic location collection

Self-reported state of birth survey data was available for 47, 473 23andMe customers. However, since overlap of these customers with our cohorts was poor, we also chose to include data from a question on current zip code of residence. This provided an additional 34, 351 zip codes of current residence. In cases where both the zip code of residence and state of birth were available, we used state of birth information. To obtain state information from zip codes, we translated zip codes to their state locations using a zip code database available from (http://federalgovernmentzipcodes.us/).

In total, we had 50, 697 individuals with available location information. About one third of each of our cohorts had location information; 1, 970 African Americans, 2, 944 Latinos, and 45, 783 European Americans were used in our geographic analyses.

### Ancestry estimation method

#### Ancestry Composition

We apply *Ancestry Composition*, a three-step pipeline that efficiently and accurately identifies the ancestral origin of chromosomal segments in admixed individuals, which is described in (Durand *et al.*, in prep). We apply the method to genotype data that have been phased using a reimplementation of *Beagle*^75^. Ancestry composition applies a string kernel support vector machines classifier to assign ancestry labels to short local phased genomic regions, which are processed using an autoregressive pair hidden Markov model to simultaneously correct phasing errors and produce reconciled local ancestry estimates and confidence scores based on the initial assignment. Lastly, these confidence estimates are recalibrated using isotonic regression models. This results in both precision and recall estimates that are greater than 0.90 across many populations, and on a continental level, have rates of 0.982–0.994 for precision, and recall rates of 0.935–0.993, depending on populations (see Table 1, Durand *et al.* in prep, and Figure S15). We note that here, and throughout the manuscript, African ancestry corresponds to Sub-Saharan African ancestry (including West African, East African, Central and South African populations, but excluding North African populations from the reference set). For more details on the the ancestry estimation method, referred to as *Ancestry Composition*, see Durand *et al.*, in prep.

#### Aggregating local ancestry information

23andMe’s *Ancestry Composition* method provides estimates of ancestry proportions for several world-wide populations at each window of the genome. To estimate genome-wide ancestry proportions of European, African, and Native American ancestry, we aggregate over populations to estimate the total likelihood of each population, and using a majority threshold of 0.51, if any window has a majority of a continental ancestry, we include it in the calculation of genome-wide ancestry, which is estimated as the number of windows passing the threshold for each ancestry over the total number of windows. Since some windows may not pass our threshold for any population, they remain unassigned, making it possible for estimates for all ancestries to not sum to 100%, resulting in population averages that likewise may not sum to 100%. We allow for this unspecified ancestry to reduce the error rates of our assignments, so, in some sense, our estimates may be viewed as lower bounds on ancestry, and it is possible that individuals carry more ancestry than estimated. In practice, we typically assign nearly all windows, with an average of about 1–2% unassigned ancestry, so we do not expect it to affect our results, with the exception of Native American ancestry, which we discuss below.

### Validation of non-European ancestry in African Americans and European Americans

Although our *Ancestry Composition* estimates are well calibrated and have been shown to accurately estimate African, European, and Native American ancestry in tests of precision and recall (see Durand *et al.*, in prep), we were concerned that low levels of non-European ancestry in European Americans that we detected may represent an artifact of *Ancestry Composition*. Hence, we pursued several lines of investigation to provide evidence that estimates of African and Native American ancestry in European Americans are robust and not artifacts.

African ancestry in European Americans is not likely to be driven by survey errors as the number of European Americans with African ancestry is 10 times larger than our estimates of survey errors. Furthermore, the ancestry profiles of self-reported European Americans with African ancestry are distinct from all other cohorts: their African ancestry is much lower than for a random sample of African Americans, and the majority of these individuals do not carry any appreciable amount of Native American ancestry, distinguishing their ancestry profiles from Latinos (see Figure S1C). A potential source of bias in our estimates is from errors in the ancestry inference algorithm. To show that our estimates are not the result of ancestry composition errors or biases, we validated the estimates of low levels of African ancestry in European Americans using an independent admixture inference method that is not model-based^76^. We also compared to previously published 1000 Genomes Project consensus estimates^77^. Another line of evidence supporting the detected ancestry in European Americans in the US is that we observe a substantially lower occurence of Native American and African ancestry in individuals who self-report four grandparents born in the same European country. Lastly, the segments of the inferred African and Native American are uniformly distributed across the genome. The segment lengths follow a similar distribution to segments in Latinos and African Americans, and using our admixture dating, we find that these introgressed segments likely arose from admixture within the last 20 generations, which liekly took place within the Americas. These lines of evidence suggest that Native American and African segments represent true segments of Native American and African introgression that were introduced after the transcontinental migrations beginning in the 1500’s.

#### Well calibrated and consistent estimates

First, we note that estimates from our ancestry composition are extremely well calibrated, with correlations of African, European, and Native American ancestry estimates showing *r*^2^ > 0.98 on 1000 Genomes African American and Latino population samples (Figure S15). Comparisons of our estimates with those published by the 1000 Genomes Consortium show the high consistency across populations and individuals. We note that our estimates of Native American ancestry are conservative. Indeed, when our *Ancestry Composition* assignment probabilities do not pass over the confidence threshold, including signals of Native American ancestry together with general East Asian/Native American ancestry (but not East Asian) recapitulates estimates from the 1000 Genomes Project consensus estimates. We note that five individuals from the ASW population from the 1000 Genomes Project have poor consistency in their estimates. These individuals have a large amount of Native American ancestry that was not modeled by the 1000 Genomes Project estimates. That these particular individuals were sampled in Oklahoma, and carry significant Native American ancestry, is supported by our own high estimates of Native American ancestry in 23andMe self-reported African Americans from Oklahoma.

#### Low detection of African and Native American ancestry in Europeans

We looked at whether all individuals who are expected to carry solely European ancestry also have simliar rates of detection of non-European ancestry. To this end, we generated a cohort of 15, 289 23andMe customers who reported that all four of their grandparents were born in the same European country. The use of four grandparent birth-country has been utilized as a proxy for assessing ancestry, see for example^78, 79^. We then examined ancestry composition results for these individuals, and calculated how at what rate we detected at least 1% African and at least 1% Native American ancestry. Overall, we find very low levels of African and Native ancestry, with 0.98% of Europeans showing African ancestry, and 0.26% of Europeans carry Native American ancestry. These levels are substantially lower than the 3.5% and 2.7% of European Americans that carry African and Native American ancestry, respectively. Furthermore, most European countries exhibit no individuals with substantial non-European ancestry, and the presence of individuals with African and Native American ancestry is limited to countries that had major ports in the Atlantic trade, and were known to have been highly connected to the Trans-Atlantic slave trade. Indeed, African ancestry in individuals from Europe is not unexpected, as approximately 9, 000 Africans were brought to Europe between 1501 and 1867^80^. Excluding countries that had major and minor ports in the Atlantic with strong connections to the slave trade (namely, Portugal, Spain, France, United Kingdom) and Malta, which has been the site of migrations from Africa and the Middle East (from Phoenicia and Carthage) and served as a major port since the completion of the Suez Canal. Excluding these countries, we obtain a dataset of 9, 701 Europeans, where we find African and Native American ancestry is virtually absent, with only 0.04% of individuals carrying 1% or more African ancestry, and 0.01% carrying 1% or more Native American ancestry, within the margins of survey error estimates.

#### Independent validation of African ancestry in European Americans using *f*4 statistics

We used f4 statistics from the ADMIXTOOLS software package and confirm the presence of African ancestry^76^. We used the f4 ratio test, designed to estimate the proportion of admixture from a related ancestral population, to compare admixture in European Americans versus reference European individuals. We tested whether European Americans with estimated African ancestry showed any admixture from Africans using our cohorts of individuals with estimated African ancestry and reference populations from the HGDP dataset. Admixture would be expected to result in estimates of *α* significantly different from 1. In our case, we both obtain significant estimates less than 1, and *α* values are correlated with our estimates of African ancestry. For populations used and estimates of admixture, see Table S6.

#### Detection of Native American mtDNA in European Americans and African Americans

The mitochondrial DNA (mtDNA) haplogroups A2, B2, B4b, C1b, C1c, C1d, and D1 are most prevalently found in the Americas, and which are likely to be Native American specific haplogroups as they are rarely found outside of the Americas. We assessed the fraction of individuals that carry these haplogroups to validate the likelihood of Native American ancestry in European Americans and African Americans, and show that these haplogroups are virtually absent in European controls. Since mtDNA haplogroups are assigned by classification using SNPs that segregate on these lineages, these orthogonal results provide an independent line of support for our estimates Native American ancestry in European Americans and African Americans. We find that the frequency of Native American haplogroups correlates with our estimates of genome-wide ancestry in European Americans and African Americans, and are found at appreciable fractions of individuals who are estimated to carry Native American ancestry. The frequencies of haplogroups are shown in Table S7. Furthermore, these haplogroups are virtually absent in individuals with four grandparents from a European country (21 individuals out of 15,651). Furthermore, the majority of these haplogroups are found in individuals from Spain, suggesting the possibility of gene flow returning from the Americas into Spain, which may also be reponsible for driving higher levels of genetic diversity in Europeans from the Iberian peninsula^81^. Excluding Spain, overall Native American specific haplogroups are detected in less than 0.05% of individuals with four grandparents from Europe, consistent with survey errors in reporting all four grandparent birthplaces.

#### Uniform distribution of ancestry segments

Regions of the genome that have structural variation, or show strong linkage disequilibrium (LD), have been shown to confound both admixture mapping and also influence the detection of population substructure in studies of PCA^82, 79,^ ^78^. If such regions were to drive artifacts of spurious ancestry, we would expect that segments of local ancestry would likely occur around these regions, rather than in a uniform distribution across the genome. To this end, we examined the starting positions of all African and Native American ancestry segments in European Americans and Native American ancestry in African Americans and show that segments of non-European ancestry start uniformly across the genome (see Figure S16). Although some regions, including the HLA region containing the MHC complex on chromsome 6, show higher ancestry switches reflecting difficulties in assignment due to genetic diversity (as likewise seen in African Americans and Latinos Figures S17 and S18), the majority of segments of ancestry appear uniformly distributed across the genome. Only 4% of all segment starts of African ancestry lie within the HLA region, and only about 1.4% of Native American segment starts lie in the HLA region. Thus, while we expect that some of the inferred ancestry may arise from difficulties in assigning ancestry in complex regions of the genome, only a small fraction of the estimated African and Native American ancestry in European Americans may be explained through such biases, and is not expected to give rise to significant, over 1% ancestry from any population.

#### Recent admixture dates suggest post-colonial admixture

Lastly, our recent dates for admixture suggest introgression likely ocurred in the Americas within the last 500 years. Hence, our estimates do not support that the African ancestry in European Americans stems from ancient population events that predate the migrations to the Americas. (For example, gene flow from Africa coinciding with the Moor invasion of the Mediterranean may have introduced African ancestry into the ancestral population of some European Americans.) Though such ancient events would likely not lead to African ancestry since our supervised learning algorithm would apply a European label to such segments, it is possible that European population substructure could lead to inferred segments of African ancestry in some European Americans that derive from older historical admixture events, which are not seen in modern Europeans. However, these events would lead to admixture or introgression of segments several hundred or thousand years old, and, our admixture dates for both Native American ancestry and African ancestry point to gene flow within the last 20 generations, and is not consistent with any known historical migrations within Europe during this time period.

### Ancestry analyses

#### Lower estimates of African ancestry in 23andMe African Americans

Unlike previous estimates of the mean proportion of African ancestry, which typically have ranged from 77% to 93% African ancestry^83–90, 3–5, 91–97^, our estimates, depending on exclusions, are 73% or 75%. There are several possible explanations for our low mean African ancestry. If our ancestry composition estimates are were downward biased, then the African Americans may have similar levels of African ancestry consistent with other studies, and our results are simply underestimates. However, our ancestry composition estimates are extremely well calibrated for African Americans from the 1000 Genomes Project and their consensus estimates, and see no evidence of a downward bias (see Durand *et al.*, in prep, and Figure S15).

The mean ancestry proportion of 23andMe self-reported African American is about 73%. A small fraction, about 2%, of African Americans, carry less than two percent African ancestry, which is far less than typically seen in most African Americans (Figure S14A). Further investigation reveals that the majority of these individuals (88%) have predominantly European ancestry, and others carry East Asian, South Asian and Southeast Asian ancestry, roughly in proportion to the frequencies found in the 23andMe database overall. Given the large number of non-African American individuals in the 23andMe database, even an exceeding low survey error rate of 0.02% could be sufficient to account for the number of outlier individuals we detect. Hence, we posit that these individuals represent survey errors rather than true self-reported African Americans. Exclusion of these 108 self-reported African Americans with less than 2% African ancestry from mean ancestry calculations results in a moderate rise, to 74.8%, of the mean proportion of African ancestry in African Americans.

To quantify differences in African ancestry driving mean state differences, we examined the distributions of estimates of African ancestry in African Americans from the District of Columbia (D.C.) and Georgia, that had at least 50 individuals with the lowest and highest mean African ancestry proportions (Figure S1E). We find a qualitiative shift in the two distributions of African ancestry, with D.C. showing a reduced mode, higher variance, as well as a heavier lower tail of African ancestry, corresponding to more African Americans with below-average ancestry than Georgia. Qualitative differences in the distributions of African ancestry proportions in African Americans from states with higher and lower mean ancestry appear to be driven by both a shift in the mode of the distribution as well as a heavier left tail reflecting more individuals with a minority of African ancestry (Figure S1). We posit that differences among states could be due to differences in admixture, differences in self-identity, or differences in patterns of assortative mating, whereby individuals with similar ancestry may preferentially mate. For example, greater levels of admixture with Europeans would both shift the mode and result in more African American individuals who have a minority of African ancestry. Alternatively, a shift toward African American self-identity for individuals with a majority of European ancestry (possibly due to changes in cultural or social forces) would likewise result in lower estimates of mean African ancestry. Lastly, assortative mating would work to maintain or increase the variance in ancestry proportions, though assortative mating alone could not shift the mean proportion of African ancestry in a population.

#### Sex bias in ancestry contributions

Sex bias in ancestry contributions, often assessed through ancestry of mtDNA and Y chromosome haplogroups, is also manifested in unequal estimates of ancestry proportions on the X chromosome, which has an inheritance pattern that differs between males and females. The X chromosome more closely follows female ancestry contributions since males contribute half as many X chromosomes. Comparing ancestry on the X chromosome to the autosomal ancestry allows us to infer whether that ancestry historically entered via males (lower X ancestry) or by females (higher X ancestry). Under equal ancestral contributions from both males and females, the X chromosome should show the same levels of admixture as the genome-wide estimates. To look for evidence of unequal male and female ancestry contributions in our cohorts, we examined ancestry on the X chromosome (NRY region), which follows a different pattern of inheritance from the autosomes. In particular, estimates of ancestry on the X chromosome have been shown to have higher African ancestry in African Americans^10^. We calculate ancestry on the X chromosome as the estimate of ancestry on just windows on the X, and compare to genome-wide estimates (which do themselves include the X chromosome). It should be noted that these calculations differ among males and females, since the X chromosome is diploid in females and thus has twice as many windows in calculation of genome-wide mean proportions. However, our results still allow a peek into sex bias as the overall contribution of the X chromosome to the genome-wide estimates is small. We note that since our ancestry estimation method conservatively assigns Native American ancestry, we expect that much of the remaining unassigned ancestry may be due to Native American ancestry assigned as broadly East Asian/Native American, which is not included in these values (see Durand *et al.* in prep, Table 1, Figure 1, reprinted here as Figure S15).

To infer estimates of male and female contributions from each ancestral population, we estimated the male and female fractions of ancestry that total the genome-wide estimates and which minimize the mean square error of the X chromosome ancestry estimates. We assume that overall male and female contributions are each 50% (∑_*pop*_ *f*_*pop,male*_ = 0.5 and ∑_*pop*_ *f*_*pop,female*_ = 0.5). We assume that the total contribution from males and females of a population gives rise to the autosomal ancestry fraction (*f*_*pop,male*_ + *f*_*pop,female*_ = *auto*_*pop*_). We then compute, using a grid search, the predicted X chromosome estimates from *f*_*pop,male*_,*f*_*pop,female*_ for each *pop* Є {*African, NativeAmerican, European*}, which are calculated, like in^7^, as

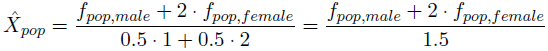

We choose the parameters of male and female contributions that minimize the mean squared error of the X ancestry estimates and the predicted 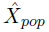. These are the estimates of male and female ancestry fractions under a single simplistic population mixture event, that best fit our X chromosome ancestry estimates observed.

For African Americans, we obtain the best fit estimates of:

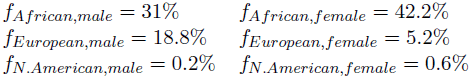

Giving rise to a male:female European ratio of 3.6, meaning that of European ancestors to African Americans, over three times as many were male as were female.

For Latinos, we obtain the following estimates of ancestry fractions from males and females:

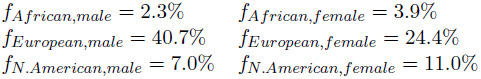

For European Americans, we obtain the following estimates:

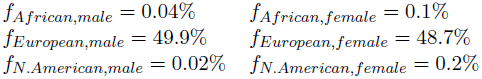

#### Population density correlations

From the 2010 Census Brief on the black population^35^, we calculated the correlation between the number of reported African Americans living in a state, and our sample of African Americans from that state. The correlation is strong, with p-value of 9.5 × 10^−14^, suggesting that our low sample sizes from states in the Mountain regions in the US is expected from estimates of population density.

African ancestry in European Americans most frequently occurs in individuals from states with high African American population density, and is rare in states with few African Americans. This observation led us to look at the correlation between population density (as a percent of state population using self-reported ethnicity from the 2010 US Census) and state mean levels of ancestry.

To examine the interaction between population density of minorities and ancestry, we used the 2010 US Census demographic survey by state^1^ (http://www.census.gov/2010census/). We compare the state population density to the mean estimated admixture proportion of individuals from that state, fitting linear regressions, and generating figures using `geom smooth(method = "lm", formula = y ~ x)` from the ggplot2 package in R (www.R-project.org).

We find that African ancestry in European Americans is strongly correlated with the population density of African Americans in each state. We find that the higher the state density proportion of African Americans, the more African ancestry is found in European Amerians from that state, reflecting the complex interaction of genetic ancestry, historical admixture, culture, and self-identified ancestry.

#### Generating the distribution of ancestry tracts

We generate ancestry segments as defined as continuous blocks of ancestry, estimating the best guess of ancestry at each window to define segments of each ancestry. Assigning the most likely ancestry at each window results in fewer spurious ancestry breaks and allows for a smaller upward bias in admixture dates, since breaks in ancestry segments push estimates of dates further back in time. We measure segment lengths using genetic distances, by mapping segment start and end physical positions to the HapMap genetic map.

#### Admixture dating

To estimate the timeframe of admixture events, we test a simple two-event, three-population admixture model using *TRACTS*^44^. We use a grid-search optimization to find four optimal parameters for the times of two admixture events and the proportions of admixture. We are limited to simple admixture models due to the computationally intensive grid search, as we were unable to obtain likelihoood convergence using any of the built-in optimizers. The model tested is as follows: two populations admix *t*1 generations ago, with proportion *frac*1 and 1 − *frac*1 respectively. A third population later mixes in *t*2 generations ago, with proportion *frac*2.

Both our ancestry segments and prior results supported a model with an earlier date of Native American admixture^44, 26^. We estimated likelihoods over plausible grid of admixture times and fractions for African Americans, Latinos, and European Americans to estimate dates of initial Native American and European admixture and subsequent African admixture. For European Americans, we find the best fit of Native/European admixture at about 12 generations ago, with African admixture at 4 generations ago. In African Americans, our dates are similar, with initial admixture 12 generations ago, and African admixture 6 generations ago, consistent with other admixture inference methods dating African American admixture. Lastly, our grid search finds dates for Latino admixture suggesting Native/European occurred about 11 generations ago and later African admixture 7 generations ago.

#### Logistic regression modeling of self-identity

We examine the probabilistic relationship between self-identity and genetically inferred ancestry. To explore the interaction between genetic ancestry and self-reported identity, we estimated the proportion of individuals that identify as African American and European American, partitioned by levels of African ancestry (Figure 5A). Jointly considering the cohorts of European Americans and African Americans, we examined the relationship between an individual’s genome-wide African ancestry proportion and whether they self-report as European American or African American. We note a strong dependence on the amount of African ancestry, with individuals carrying less than 20% African ancestry identifying largely as European American, and those with greater than 50% reporting as African American (Figure 4). To test the significance of this relationship, we fit a logisitic regression model, using python’s `statsmodels` package, predicting self-reported ancestry using proportion African ancestry, sex, age, intercept, and interaction variables. For a full characterization of terms and logistic models, see Supplemental Table S5 and Figure S13. We find that significant covariates include proportion African ancestry with a p-value of < 0.001, age×ancestry interaction and sex×ancestry interaction. Sex as a predictor of self-reported ancestry is not significant, with a coefficient of 0.29 and p-value 0.084. The proportion of African ancestry predicts self-reported ancestry significantly, with a coefficient of 20.1 (95% CI: 18.0–22.2).

## Acknowledgements

We thank the customers of 23andMe who answered surveys and participated in this research. We are grateful to Dr. Jeffrey C. Long at the University of New Mexico, Dr. Claudio Saunt at the University of Georgia, Sarah Abel at Centre International des Recherches sur les Esclavages, CNRS, Paris for invaluable discussions and comments on a manuscript draft. We thank Nick Patterson and Priya Moorjani for helpful statistical discussions on *f* statistics. Of course, all mistakes and inaccuracies are our own.

Thanks to all the employees of 23andMe, who together have made this research possible, especially Emma Pierson, and Scott Hadly.

KB, ED, and JLM are current employees of 23andMe, Inc, while JMM is a former employee, and have private equity interest. This work was supported by NIH award 2R44HG00698102.

