## Supplemental Tables and Figures for "The genetic ancestry of African, Latino, and European Americans across the United States"

Table S1: **Introduction text to the ethnicity survey.** We note that the text clearly states that the survey will be used in ancestry-related research.

Medical researchers in the United States regularly assess research participants race and ethnicity to ensure that inclusion in their research is fair and equitable. The definitions of race and ethnicity used in research are the same as the US Census categories, meaning medical researchers define race and ethnicity socially rather than biologically.

If you were born or live in the United States, this survey seeks to understand how you identify yourself in terms of these socially-defined categories. Whether or not you are from the United States, the survey asks about your geographic roots.

Your answers will help 23andMe understand the genetic diversity of these categories. Your responses to this survey may be used in both health-related and ancestry-related research and in summarizing the ethnic breakdown of participants for some of our federally-funded studies, such as those funded by the National Institutes of Health (NIH). Over time your responses will enable 23andMe to improve its health and ancestry reports. Want to help improve 23andMes health and ancestry features? Tell us how you identify in terms of ethnic and racial categories. Tell us what you know about your geographic roots. Your answers may lead not only to new research findings, but also to new 23andMe health and ancestry reports. This survey is about how you identify yourself in terms of the socially-defined categories of ethnicity and race, and about your geographic roots. Your responses to this survey may be used in both health-related and ancestry-related research, and in summarizing the ethnic breakdown of participants for some of our federally-funded studies, such as those funded by the National Institutes of Health (NIH).

Estimated time to complete: Less than 5 minutes

Table S2: **Mean ancestry proportions and sample sizes of 23andMe African Americans, European Americans, and Latinos.** To protect participant privacy, ancestries have been rounded, and states with fewer than 10 individuals from a cohort are not reported. Sample sizes between 10 and 49 individuals are denoted by (*), between 50 and 99 individuals by (**) and 100 or more individuals as (***). Mean levels of European (Eur.), African (Af.) and Native American (N. Am.) ancestry are reported for each state.

| | African Americans | | | | European Americans | | | | Latinos | | | |
|---|---|---|---|---|---|---|---|---|---|---|---|---|
| State | Af. | N.Am. | Eur. | Size | Af. | N.Am. | Eur. | Size | Af. | N.Am. | Eur. | Size |
| Alabama | 81% | 0.7% | 17% | * | 0.5% | 0.1% | 98.9% | *** | - | - | - | - |
| Alaska | - | - | - | - | 0.2% | 0.4% | 98.5% | *** | - | - | - | - |
| Arizona | - | - | - | - | 0.1% | 0.1% | 99.2% | *** | 4% | 18% | 69% | ** |
| Arkansas | 80% | 0.5% | 18% | * | 0.2% | 0.1% | 99.3% | *** | 5% | 10% | 80% | * |
| California | 73% | 0.8% | 24% | *** | 0.2% | 0.3% | 98.1% | *** | 4% | 19% | 65% | *** |
| Colorado | 72% | 0.8% | 25% | * | 0.2% | 0.1% | 99.2% | *** | 4% | 18% | 67% | ** |
| Connecticut | 77% | 0.5% | 21% | * | 0.1% | 0.0% | 99.0% | *** | 10% | 9% | 75% | * |
| DC | 70% | 0.6% | 28% | ** | 0.1% | 0.1% | 99.2% | *** | 14% | 9% | 64% | * |
| Delaware | - | - | - | - | 0.2% | 0.1% | 99.5% | *** | - | - | - | - |
| Florida | 81% | 0.5% | 17% | * | 0.3% | 0.1% | 98.7% | *** | 6% | 7% | 80% | *** |
| Georgia | 81% | 0.6% | 17% | ** | 0.4% | 0.1% | 99.3% | *** | 16% | 8% | 71% | * |
| Hawaii | - | - | - | - | 0.2% | 0.1% | 97.6% | *** | 4% | 5% | 57% | * |
| Idaho | - | - | - | - | 0.2% | 0.3% | 98.7% | *** | - | - | - | - |
| Illinois | 74% | 0.5% | 24% | *** | 0.1% | 0.1% | 99.1% | *** | 8% | 19% | 63% | *** |
| Indiana | 73% | 0.6% | 25% | * | 0.1% | 0.1% | 99.3% | *** | 5% | 8% | 83% | * |
| Iowa | - | - | - | - | 0.1% | 0.1% | 99.5% | *** | 6% | 5% | 79% | * |
| Kansas | 69% | 0.5% | 29% | * | 0.2% | 0.1% | 99.5% | *** | 4% | 14% | 75% | * |
| Kentucky | 69% | 0.4% | 29% | * | 0.3% | 0.1% | 99.3% | *** | 4% | 4% | 90% | * |
| Louisiana | 75% | 0.8% | 23% | ** | 0.6% | 0.3% | 98.5% | *** | 22% | 4% | 70% | * |
| Maine | - | - | - | - | 0.1% | 0.1% | 99.6% | *** | - | - | - | - |
| Maryland | 72% | 0.6% | 26% | * | 0.1% | 0.1% | 99.2% | *** | 10% | 7% | 76% | * |
| Massachusetts | 73% | 0.5% | 25% | * | 0.1% | 0.1% | 98.1% | *** | 11% | 10% | 73% | * |
| Michigan | 75% | 0.7% | 23% | ** | 0.1% | 0.1% | 98.9% | *** | 15% | 9% | 69% | * |
| Minnesota | - | - | - | - | 0.0% | 0.1% | 99.4% | *** | 12% | 10% | 70% | * |
| Mississippi | 80% | 0.6% | 18% | * | 0.3% | 0.2% | 99.1% | *** | - | - | - | - |
| Missouri | 76% | 0.5% | 22% | ** | 0.2% | 0.1% | 99.4% | *** | 14% | 8% | 76% | * |
| Montana | - | - | - | - | 0.1% | 0.3% | 99.2% | *** | - | - | - | - |
| Nebraska | - | - | - | - | 0.1% | 0.1% | 99.3% | *** | - | - | - | - |
| Nevada | - | - | - | - | 0.2% | 0.3% | 98.2% | *** | - | - | - | - |
| New Hampshire | - | - | - | - | 0.1% | 0.0% | 99.5% | *** | - | - | - | - |
| New Jersey | 72% | 1.1% | 25% | ** | 0.2% | 0.0% | 98.3% | *** | 9% | 10% | 73% | ** |
| New Mexico | - | - | - | - | 0.2% | 0.4% | 98.7% | *** | 3% | 20% | 67% | ** |
| New York | 75% | 0.9% | 22% | *** | 0.1% | 0.1% | 97.8% | *** | 15% | 8% | 69% | *** |
| North Carolina | 74% | 0.6% | 23% | ** | 0.4% | 0.1% | 98.9% | *** | 17% | 5% | 75% | * |
| North Dakota | - | - | - | - | 0.1% | 0.3% | 98.8% | *** | - | - | - | - |
| Ohio | 73% | 0.6% | 24% | ** | 0.2% | 0.0% | 99.1% | *** | 11% | 7% | 78% | * |
| Oklahoma | 73% | 0.9% | 25% | * | 0.3% | 0.2% | 99.1% | *** | 8% | 11% | 72% | * |
| Oregon | - | - | - | - | 0.2% | 0.2% | 98.8% | *** | 3% | 11% | 74% | * |
| Pennsylvania | 72% | 0.6% | 26% | *** | 0.1% | 0.0% | 99.0% | *** | 15% | 8% | 72% | ** |
| Rhode Island | - | - | - | - | 0.1% | 0.1% | 98.7% | *** | - | - | - | - |
| South Carolina | 83% | 0.7% | 15% | * | 0.5% | 0.2% | 99.0% | *** | - | - | - | - |
| South Dakota | - | - | - | - | 0.0% | 0.0% | 99.8% | *** | - | - | - | - |
| Tennessee | 77% | 0.5% | 21% | ** | 0.3% | 0.1% | 99.1% | *** | 2% | 6% | 89% | * |
| Texas | 78% | 0.7% | 20% | *** | 0.2% | 0.3% | 98.9% | *** | 5% | 21% | 64% | *** |
| Utah | - | - | - | - | 0.1% | 0.2% | 98.9% | *** | 1% | 13% | 78% | * |
| Vermont | - | - | - | - | 0.1% | 0.2% | 99.1% | *** | - | - | - | - |
| Virginia | 74% | 0.6% | 23% | ** | 0.4% | 0.1% | 98.9% | *** | 13% | 10% | 71% | * |
| Washington | 66% | 0.9% | 30% | * | 0.1% | 0.2% | 99.0% | *** | 7% | 9% | 76% | * |
| West Virginia | 64% | 0.2% | 34% | * | 0.2% | 0.1% | 98.9% | *** | - | - | - | - |
| Wisconsin | 71% | 0.5% | 27% | * | 0.1% | 0.1% | 99.4% | *** | 11% | 14% | 68% | * |
| Wyoming | - | - | - | - | 0.1% | 0.1% | 99.6% | *** | - | - | - | - |

Table S3: Mean ancestry proportions and sample sizes of 23andMe African Americans, by region. Sample sizes between 100 and 499 individuals are denoted by (*), between 500 and 999 individuals by (**) and 1000 or more individuals as (***). Mean levels of European, African and Native American ancestry are reported for each subpopulation.

| Region | African ancestry | European ancestry | Native American ancestry | Sample size | (States included in region) |
|---|---|---|---|---|---|
| West | 72.6% | 24.3% | 0.9% | * | New Mexico, Hawaii, California, Montana, Oregon, Utah, Arizona, Idaho, Nevada, Wyoming, Alaska, Washington, Colorado |
| Midwest | 73.6% | 24.1% | 0.6% | * | Missouri, Nebraska, Ohio, Kansas, Michigan, Wisconsin, Indiana, Illinois, Minnesota, Iowa, North Dakota, South Dakota |
| Northeast | 73.2% | 24.3% | 0.8% | ** | Rhode Island, Pennsylvania, Vermont, New York, New Hampshire, Massachusetts, Connecticut, New Jersey, D.C., Maine |
| South | 77.1% | 21.9% | 0.6% | ** | Alabama, Texas, Kentucky, Florida, Georgia, Virginia, Louisiana, Maryland, North Carolina, Arkansas, South Carolina, West Virginia, Oklahoma, Mississippi, Tennessee, Delaware |

Table S4: **Mean ancestry proportions and sample sizes of 23andMe Latinos by subpopulation.** Mean proportions of ancestry among Latino individuals that selected "Hispanic" who also chose to select another identity, or selected one or more other ethnicities are provided. To protect participant privacy, ancestries have been rounded to the nearest percent. Sample sizes between 100 and 499 individuals are denoted by (*), between 500 and 999 individuals by (**) and 1000 or more individuals as (***). Mean levels of European, African and Native American ancestry are reported for each subpopulation.

| Subpopulation | European | African | Native American | Sample Size |
|---|---:|---:|---:|---:|
| Central American | 53% | 9% | 26% | * |
| Mexican | 61% | 3% | 24% | *** |
| South American | 69% | 5% | 17% | ** |
| White | 73% | 5% | 14% | *** |
| Cuban | 84% | 6% | 4% | * |
| Puerto Rican | 69% | 14% | 8% | * |
| Dominican | 56% | 28% | 7% | * |
| Black | 46% | 42% | 6% | * |

Table S5: **Logistic regression model results for predicting European American versus African American self-reported identity.** Logistic regression was performed using python's module `statsmodels`. The three models shown below include the full model, a model including only the most significant parameters, and a simple model using proportion African ancestry and intercept.

Logit Regression Results

| | | | |
|---|---|---|---|
| Dep. Variable: | 0 | No. Observations: | 161460 |
| Model: | Logit | Df Residuals: | 161454 |
| Method: | MLE | Df Model: | 5 |
| Date: | Mon, 12 May 2014 | Pseudo R-squ.: | 0.9416 |
| Time: | 15:10:22 | Log-Likelihood: | -1357.8 |
| converged: | True | LL-Null: | -23269. |
| | | LLR p-value: | 0.000 |

| | coef | std err | z | P>\|z\| | [95.0% Conf. Int.] | |
|---|---|---|---|---|---|---|
| ancestry | 20.0753 | 1.069 | 18.775 | 0.000 | 17.980 | 22.171 |
| age | 4.148e-05 | 0.005 | 0.009 | 0.993 | -0.009 | 0.010 |
| sex | 0.2906 | 0.168 | 1.730 | 0.084 | -0.039 | 0.620 |
| age-ancestry-interaction | 0.0472 | 0.020 | 2.308 | 0.021 | 0.007 | 0.087 |
| sex-ancestry-interaction | -2.3224 | 0.768 | -3.024 | 0.002 | -3.828 | -0.817 |
| intercept | -7.1956 | 0.261 | -27.600 | 0.000 | -7.707 | -6.685 |

Logit Regression Results

| | | | |
|---|---|---|---|
| Dep. Variable: | 0 | No. Observations: | 161460 |
| Model: | Logit | Df Residuals: | 161456 |
| Method: | MLE | Df Model: | 3 |
| Date: | Mon, 12 May 2014 | Pseudo R-squ.: | 0.9416 |
| Time: | 15:10:25 | Log-Likelihood: | -1359.3 |
| converged: | True | LL-Null: | -23269. |
| | | LLR p-value: | 0.000 |

| | coef | std err | z | P>\|z\| | [95.0% Conf. Int.] | |
|---|---|---|---|---|---|---|
| ancestry | 19.6822 | 0.865 | 22.745 | 0.000 | 17.986 | 21.378 |
| age-ancestry-interaction | 0.0476 | 0.016 | 2.887 | 0.004 | 0.015 | 0.080 |
| sex-ancestry-interaction | -1.5871 | 0.639 | -2.485 | 0.013 | -2.839 | -0.335 |
| intercept | -7.0514 | 0.084 | -84.239 | 0.000 | -7.216 | -6.887 |

Logit Regression Results

| | | | |
|---|---|---|---|
| Dep. Variable: | 0 | No. Observations: | 161460 |
| Model: | Logit | Df Residuals: | 161458 |
| Method: | MLE | Df Model: | 1 |
| Date: | Mon, 12 May 2014 | Pseudo R-squ.: | 0.9413 |
| Time: | 15:12:43 | Log-Likelihood: | -1365.8 |
| converged: | True | LL-Null: | -23269. |
| | | LLR p-value: | 0.000 |

| | coef | std err | z | P>\|z\| | [95.0% Conf. Int.] | |
|---|---|---|---|---|---|---|
| ancestry | 21.0602 | 0.375 | 56.096 | 0.000 | 20.324 | 21.796 |
| intercept | -7.0477 | 0.084 | -84.331 | 0.000 | -7.212 | -6.884 |

Table S6: **Estimates of admixture from ADMIXTOOLS f4 test.** Estimates of admixture from Africans into European Americans, stratified by our estimates of African ancestry, are shown. Populations used for validation include 1000 Genomes populations from Italy, Great Britain, and Yoruba from Nigeria.

| X (test) | A | O (outgroup) | B (control) | C | alpha | stderr |
|---|---|---|---|---|---|---|
| European Americans 0.01-0.02 African | TSI | Chimp | GBR | YRI | 0.972757 | 0.002220 |
| European Americans >0.02 African | TSI | Chimp | GBR | YRI | 0.942362 | 0.002508 |

Table S7: **Rates of mtDNA haplogroups A, B, C and D in African Americans and European Americans with Native American ancestry**. Estimates of the number of individuals that carry Native American mtDNA haplogroups corresponds, as expected, with the estimate of genome-wide Native American ancestry. Individuals from each cohort with Native American ancestry were stratified by their estimated amount of Native American ancestry, and the number of A, B, C or D mtDNA haplogroups, and the rate of these Native American specific haplogroups is shown for each estimated amoung of Native American ancestry.

| Cohort | Prop N. Am. ancestry | N. Am. haplogroups | Total N | Rate |
|---|---|---|---|---|
| European Americans | 0.01–0.02 | 96 | 1,278 | 7.5% |
| European Americans | $> 0.02$ | 774 | 2,697 | 28.7% |
| African Americans | 0.01–0.02 | 16 | 838 | 1.9% |
| African Americans | $> 0.02$ | 34 | 305 | 11.1% |
| 4GP Europeans | all countries | 21 | 15,651 | 0.13% |
| 4GP Europeans | excl. Spain | 7 | 15,021 | 0.047% |

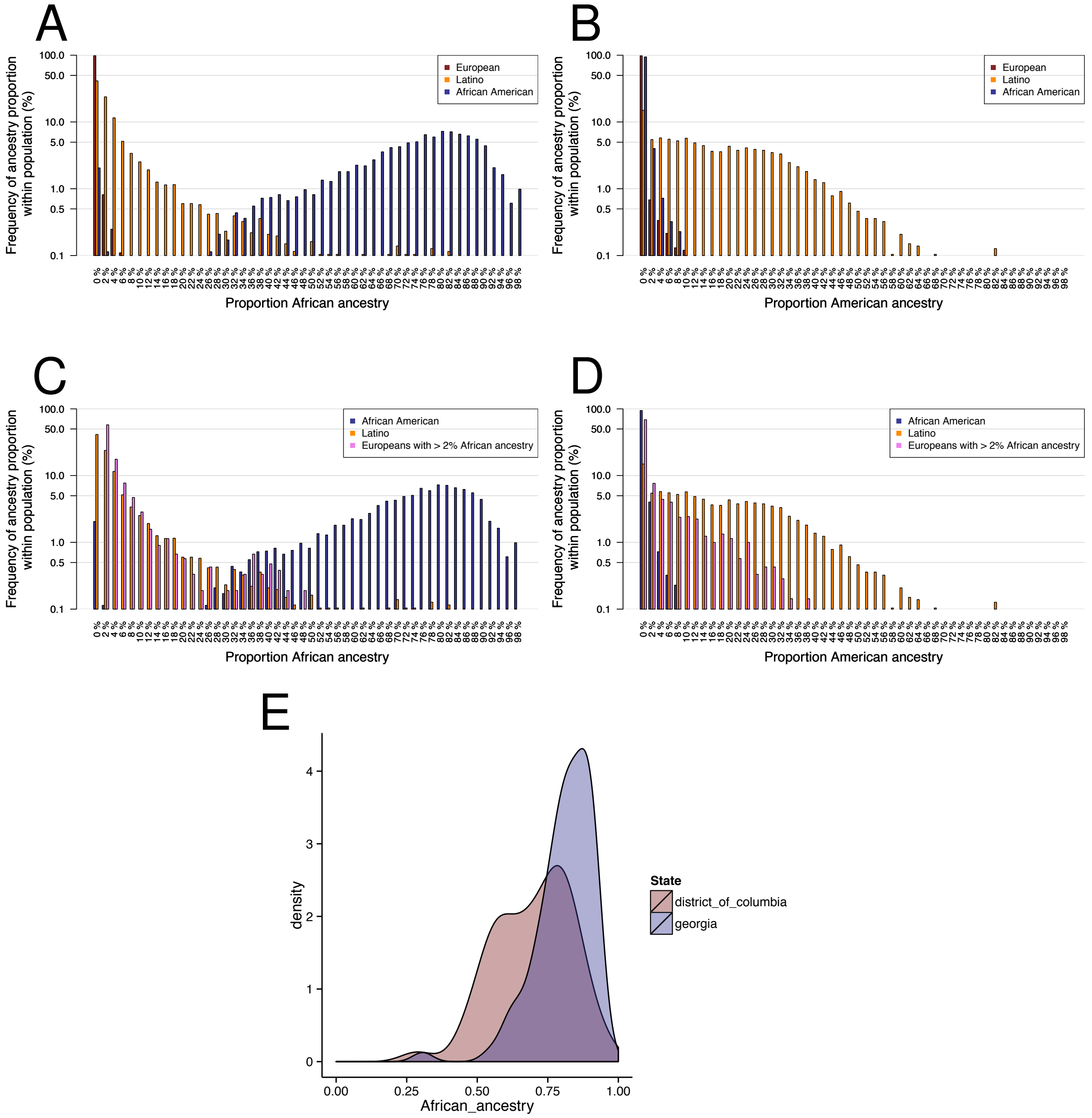


Figure S1: **Histogram of ancestry, in bins of 2%, in self-reported African American, Latino, and European American individuals.** The vertical bars represent the proportion of individuals from each self-reported cohort that are estimated to have proportion African ancestry fall within each ancestry bin. Note that the y-axis is shown in a log scale to illustrate fine-scale differences among cohorts. Histogram of African ancestry (A) and Native American (B) in European Americans (red bars), Latinos (gold bars), and African Americans (blue bars). Histogram of African (C) and Native American (D) ancestry in African Americans, Latinos, and only those European Americans that have at least 2% African ancestry. (E) Qualitative differences in African ancestry distributions in African Americans from California and Georgia. Restricted to states for which we had at least 50 individuals, D.C. had the lowest mean African ancestry, and Georgia had the highest mean African ancestry. The distribution of the ancestry proportions of self-reported African American individuals from these states are displayed using `geom_density` in `ggplot2` from `R`.

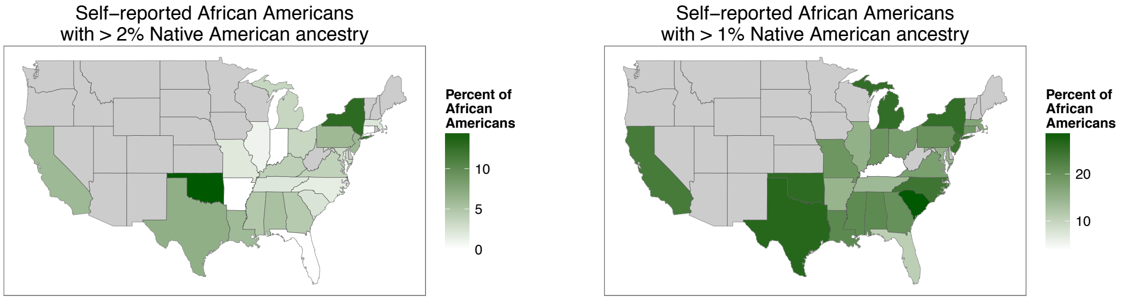


Figure S2: **Frequency of self-reported African American individuals with at least 2% (left) and 1% (right) Native American ancestry across states with at least 20 individuals.** The geographic distribution of self-reported African Americans with Native American ancestry. States with fewer than 20 individuals are excluded and shaded in gray. The proportion of individuals with Native American ancestry, out of the total number of African Americans per state, is shown by shade of green.

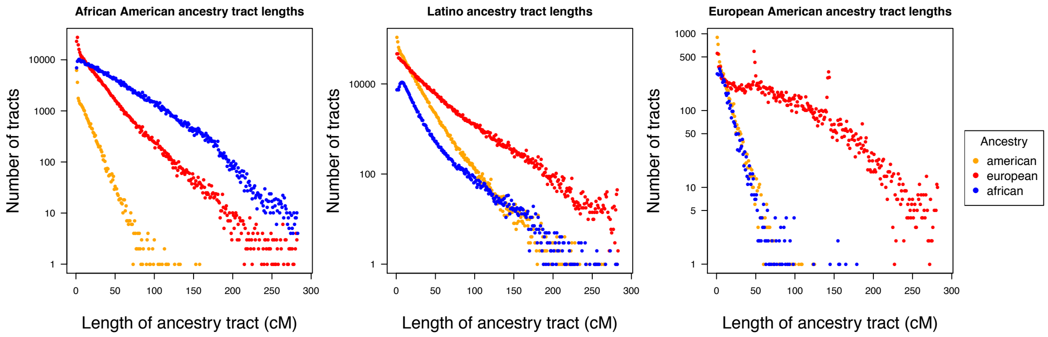


Figure S3: **Distribution of the lengths of ancestry segments for African Americans, Latinos, and Europeans with at least 2% African ancestry.** The lengths of segments, or tracts, of ancestry, and the frequency of those tracts is shown by points, colored by population. The number of ancestry tracts is shown on a log scale. Counts are shown self-reported African Americans, Latinos, and European Americans. The number of tracts of Native American (gold), European (red) and African (blue) ancestry tracts is shown for each bin of 1Mb of segment length.

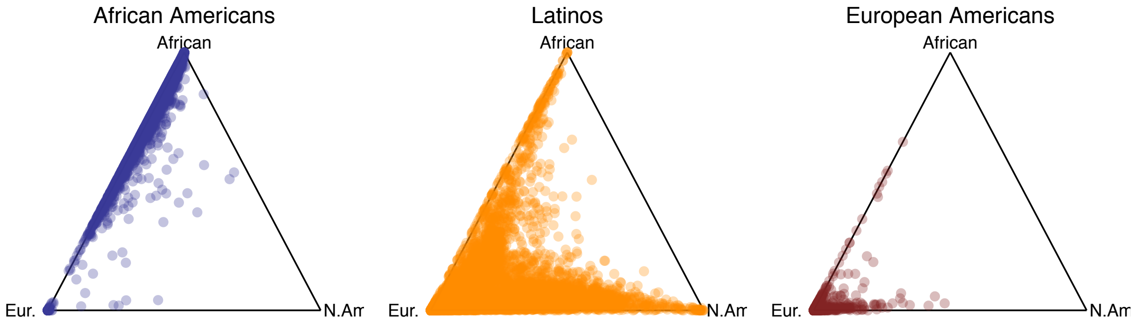


Figure S4: **Ternary plots of African, European, and Native American ancestry in self-reported African American, Latino, and European American individuals.** Each point represents a self-reported individual and is positioned within the triangle reflecting the amount of ancestry estimated from each population. Note that each individual is plotted as a semi-transparent point to convey density of individuals. Only a random sample of 10,000 of the European Americans are shown for plotting purposes.

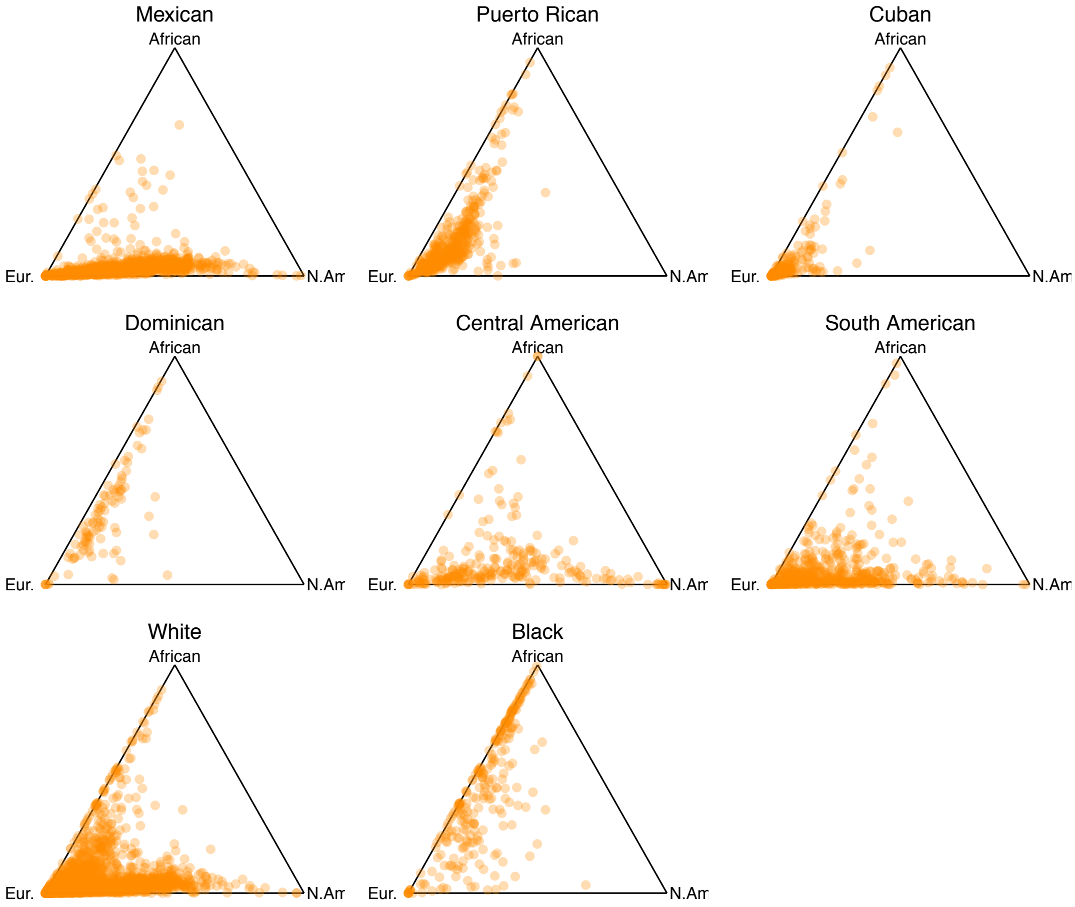


Figure S5: **Ancestry of self-reported Latinos by secondary self-reported subpopulation.** Each individual is shown projected onto the triangle by their genome-wide proportions of African, European, and Native American ancestry, by their self-reported Hispanic sub-identity. Proportion of ancestry can be computed for an individual from the distance in dropping a perpendicular line from the point to the edge opposite the vertex.

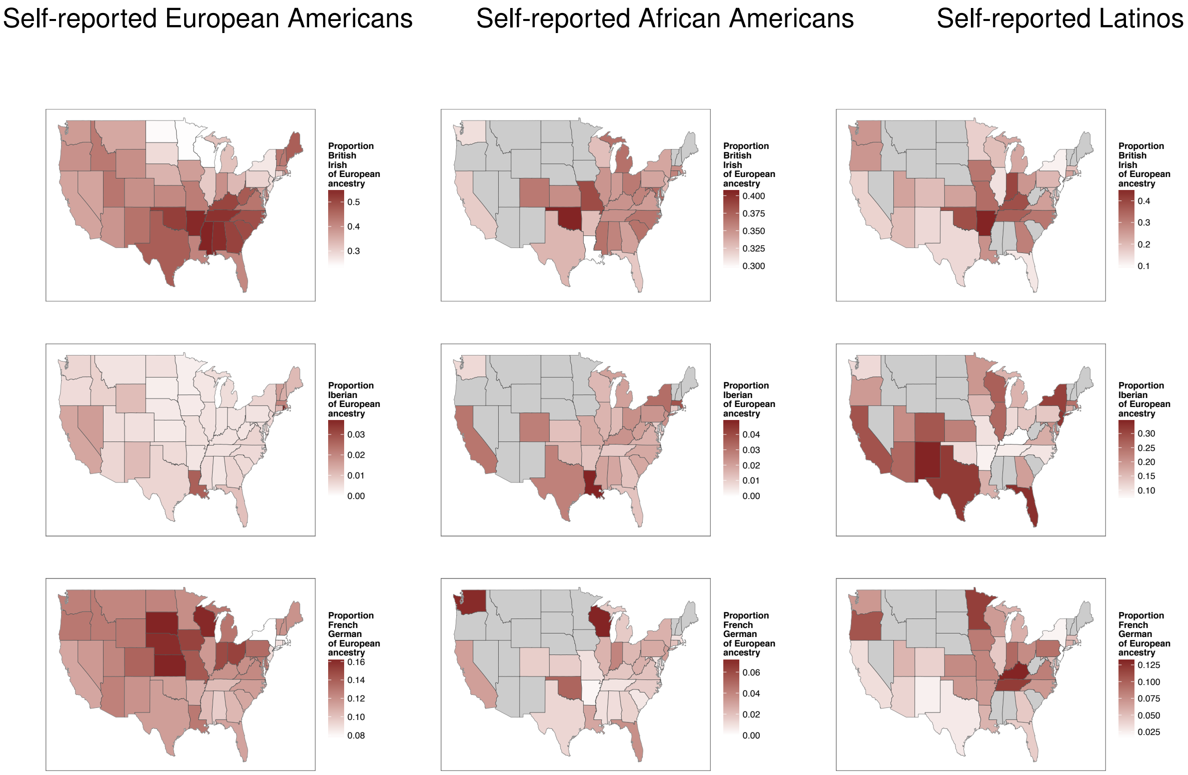


Figure S6: **Differences in the European subpopulation *Ancestry Composition* among self-reported European Americans, African Americans, and Latinos from different states.** The relative amoung of European ancestry, out of the total mean European ancestry, estimated for each state. Shown for inferred British/Irish ancestry, inferred Iberian ancestry, and inferred Italian ancestry. The proportion of sub-population ancestry, normalized by the total estimated European ancestry, for each state is shown by shade of red.

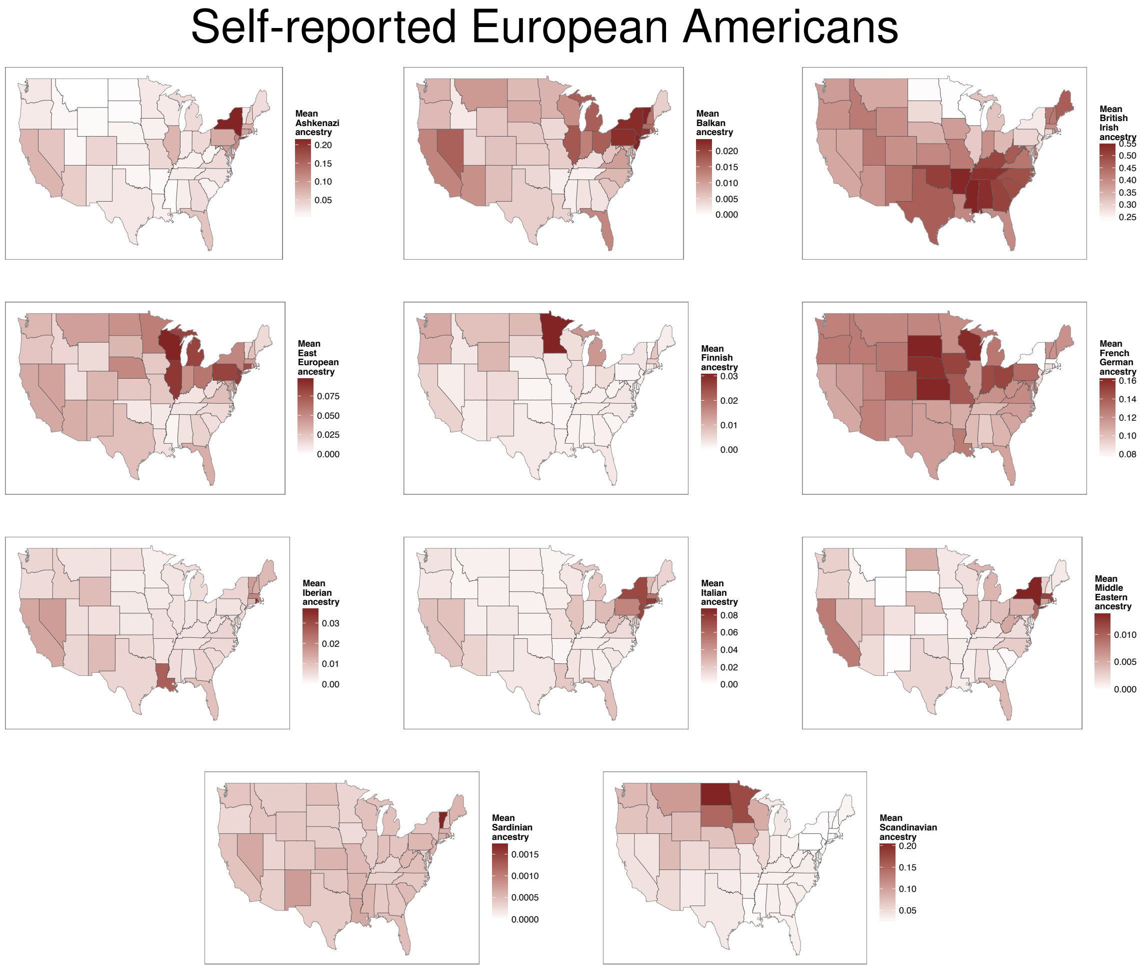


Figure S7: **Differences in the European subpopulation ancestry among self-reported European Americans from different states.** Shown for all European subpopulations that are carried at greater than 1% frequency in some state. The mean ancestry proportion among self-reported European Americans from each state is shown by shade of red. Ancestries that do not achieve at least 1% mean average ancestry in any state are not shown.

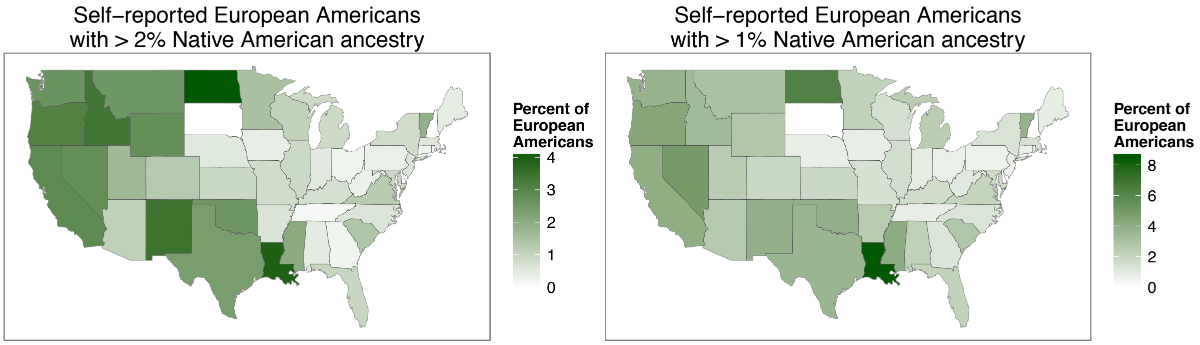


Figure S8: **Frequency of self-reported European Americans with at least 2% Native American ancestry (left) and 1% Native American ancestry (right).** The geographic distribution of self-reported European Americans with Native American ancestry. States with fewer than 20 individuals are excluded and shaded in gray. The proportion of individuals with Native American ancestry, out of the total number of European Americans per state, is shown by shade of green.

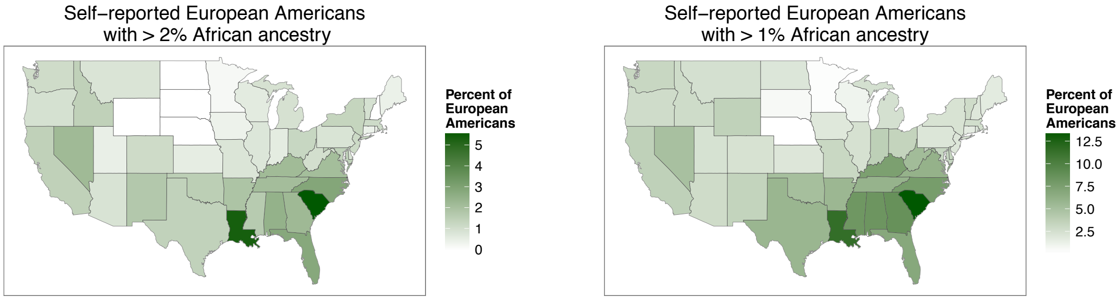


Figure S9: **Frequency of self-reported European Americans with at least 2% African ancestry (left) and 1% African ancestry (right).** The geographic distribution of self-reported European Americans with African ancestry. States with fewer than 20 individuals are excluded and shaded in gray. The proportion of individuals with African ancestry, out of the total number of European Americans per state, is shown by shade of green.

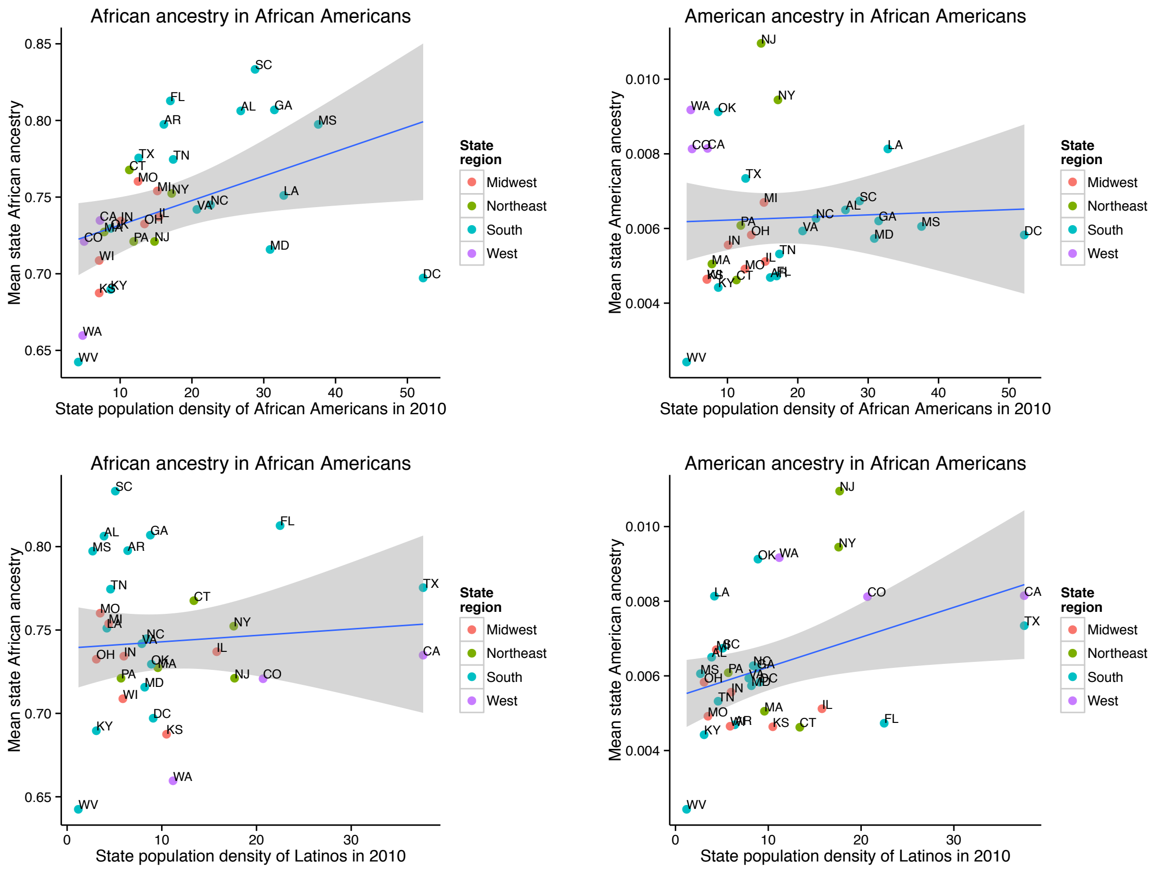


Figure S10: **Correlations of African and Native American ancestry components of African Americans with population density of African Americans and Latinos by state.** The x-axis show the state estimated population density of African Americans (top row) and Latinos (bottom row), and the y-axis show the mean state ancestry proportions. Each point represents a state and is labeled by the two-letter state abbreviation, for states with at least 10 individuals. The blue line shows a regression fit between the two variables, and the 95% confidence interval for the line fit is shown in gray. Each state is colored by region.

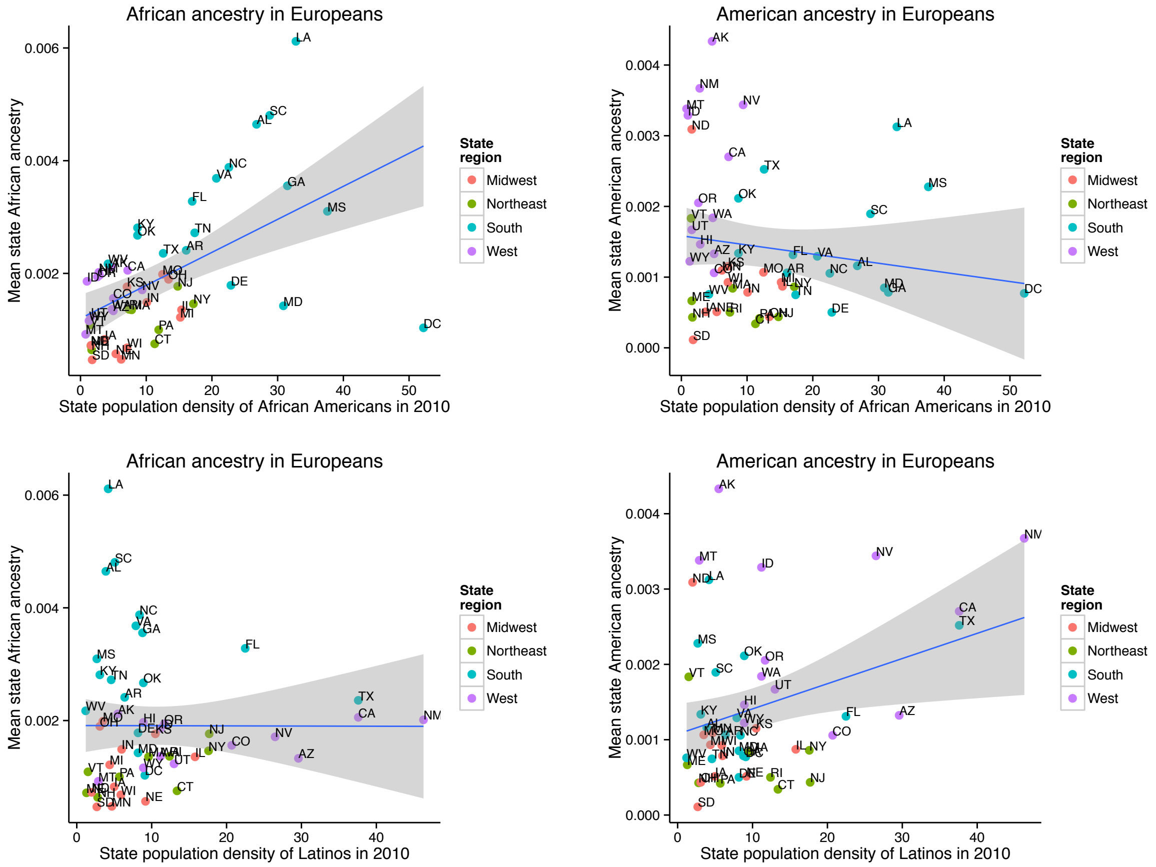


Figure S11: **Correlations of African and Native American ancestry components of European Americans with population density of African Americans and Latinos by state.**The x-axis show the state estimated population density of African Americans (top row) and Latinos (bottom row), and the y-axis show the mean state ancestry proportions. Each point represents a state and is labeled by the two-letter state abbreviation, for states with at least 10 individuals. The blue line shows a regression fit between the two variables, and the 95% confidence interval for the line fit is shown in gray. Each state is colored by region.

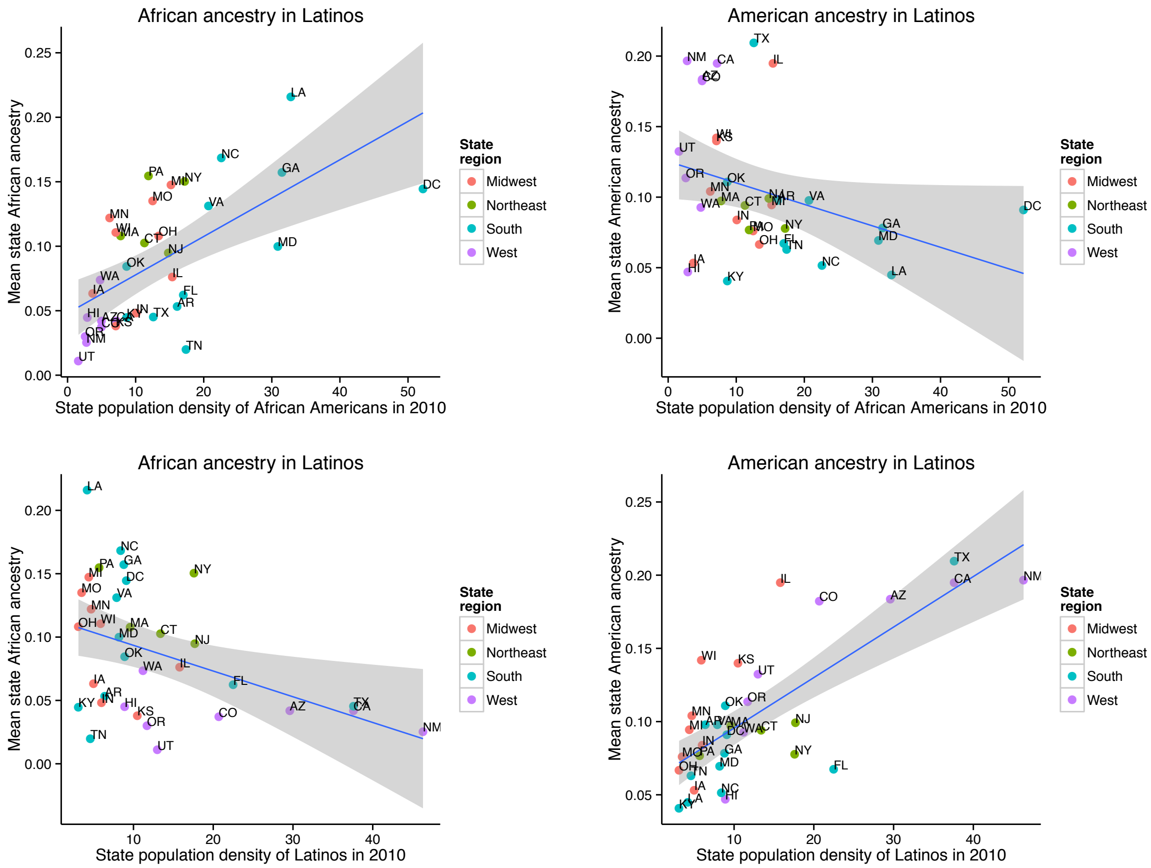


Figure S12: **Correlations of African and Native American ancestry components of Latinos with population density of African Americans and Latinos by state.**The x-axis show the state estimated population density of African Americans (top row) and Latinos (bottom row), and the y-axis show the mean state ancestry proportions. Each point represents a state and is labeled by the two-letter state abbreviation, for states with at least 10 individuals. The blue line shows a regression fit between the two variables, and the 95% confidence interval for the line fit is shown in gray. Each state is colored by region.

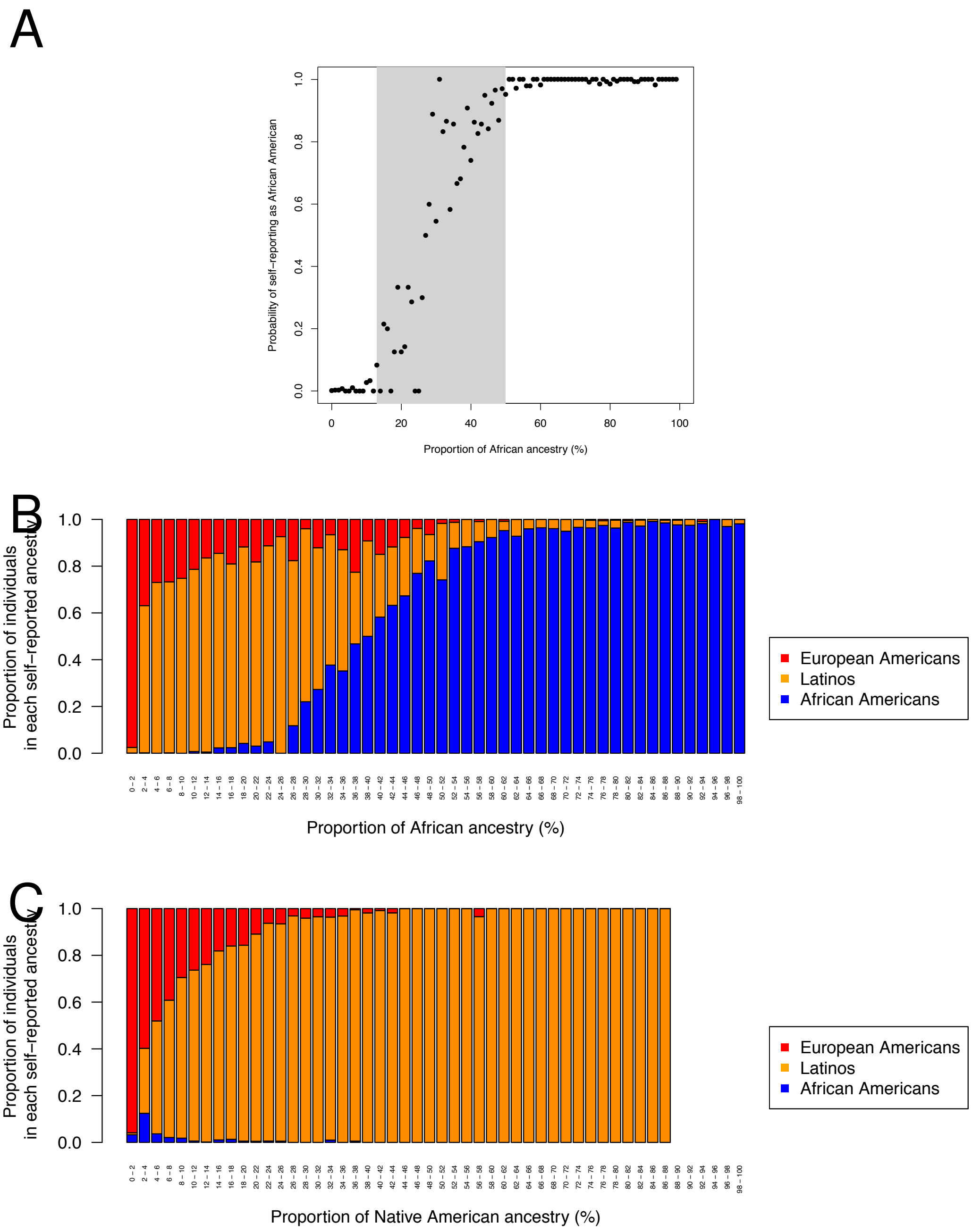


Figure S13: **Relationship between the amount of African ancestry and African American versus European American self-reported identity.** (A) Using ancestry data jointly from both African Americans and European Americans, we show the probability of self-reporting as African American by proportion of African ancestry. The probability for each bin of 1% ancestry is shown (points), and the gray area is shaded to emphasize the transition region. (B) Proportion of individuals that self-report as European American, African American, and Latino, by proportion of African ancestry. (C) The proportion of individuals that self-report as European American, African American, and Latino by the proportion of Native American ancestry.

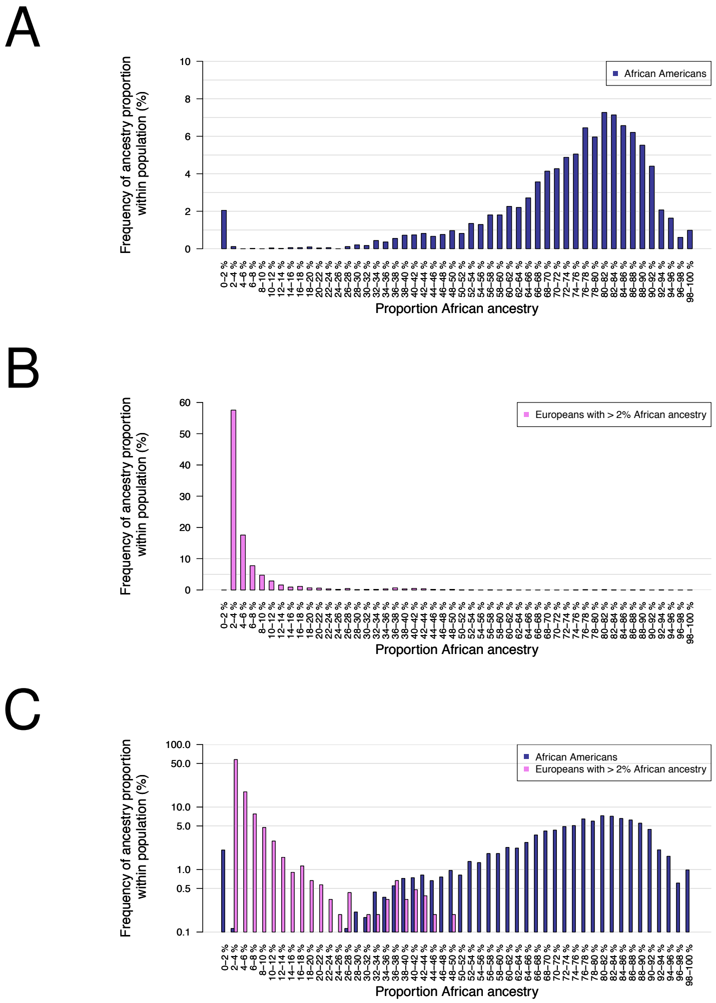


Figure S14: **Distribution of African ancestry in African Americans and European Americans.** (A) Histogram of African ancestry proportions of self-reported African Americans. (B) Histogram of those European Americans that are estimated to have at least 2% African ancestry. (C) Combined histogram of African Americans and European Americans that carry at least 2% African ancestry. Note that histogram C is shown on a log-scale to allow visualization of fine-scale differences between populations. Bins representing less than 0.1% of individuals are not shown.

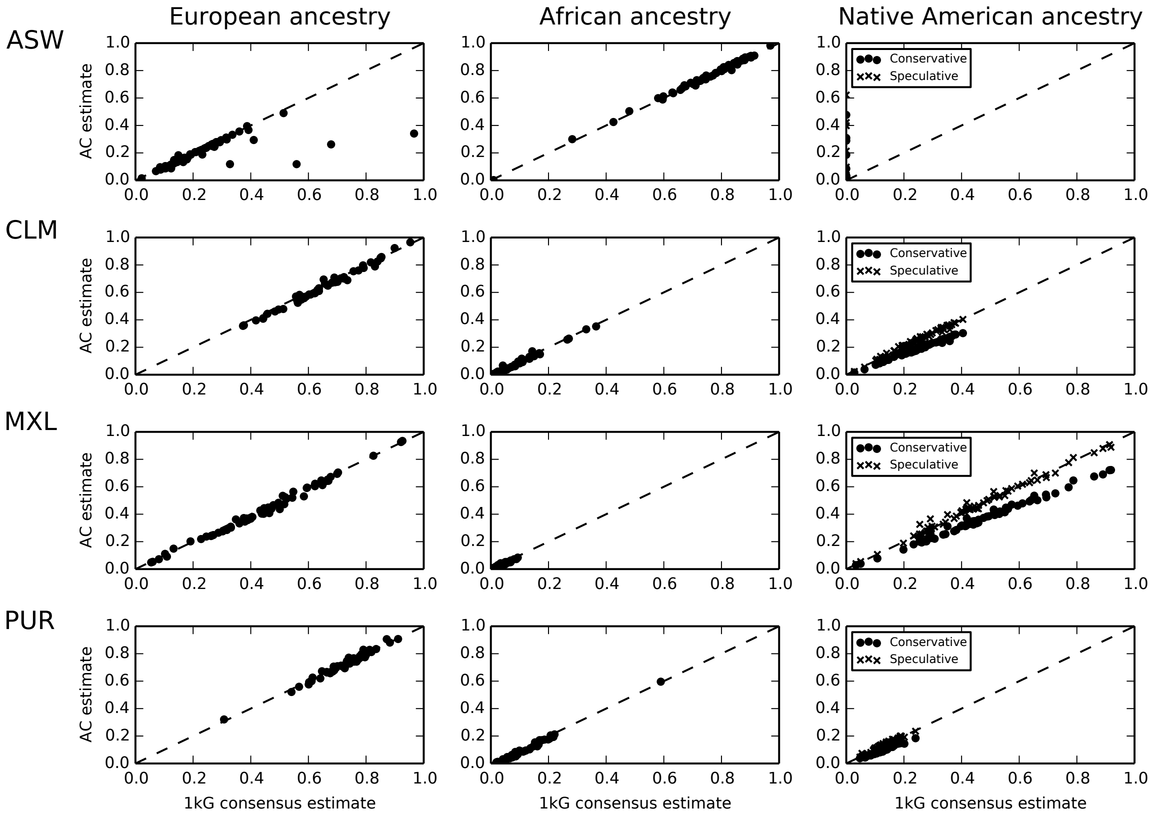


Figure S15: **Comparison of Ancestry Composition estimates with 1000 genomes consensus estimates on four recently admixed populations from the 1000 genomes project: ASW (African Americans), CLM (Colombians), MXL (Mexicans) and PUR (Puerto Ricans).** For each of the four populations, we plot the European, African and Native American admixture proportions estimated by Ancestry Composition versus the 1000 genomes consensus estimates. We note that 5 individuals from the ASW population show large amounts of Native American ancestry that was predicted as European by the 1kG consensus method. Ancestry Composition tends to underestimates the proportion of Native American ancestry in CLM, MXL and PUR compared to the 1kG consensus method (Conservative estimates), unless we allow estimates of general East Asian/Native American ancestry (Speculative estimates).

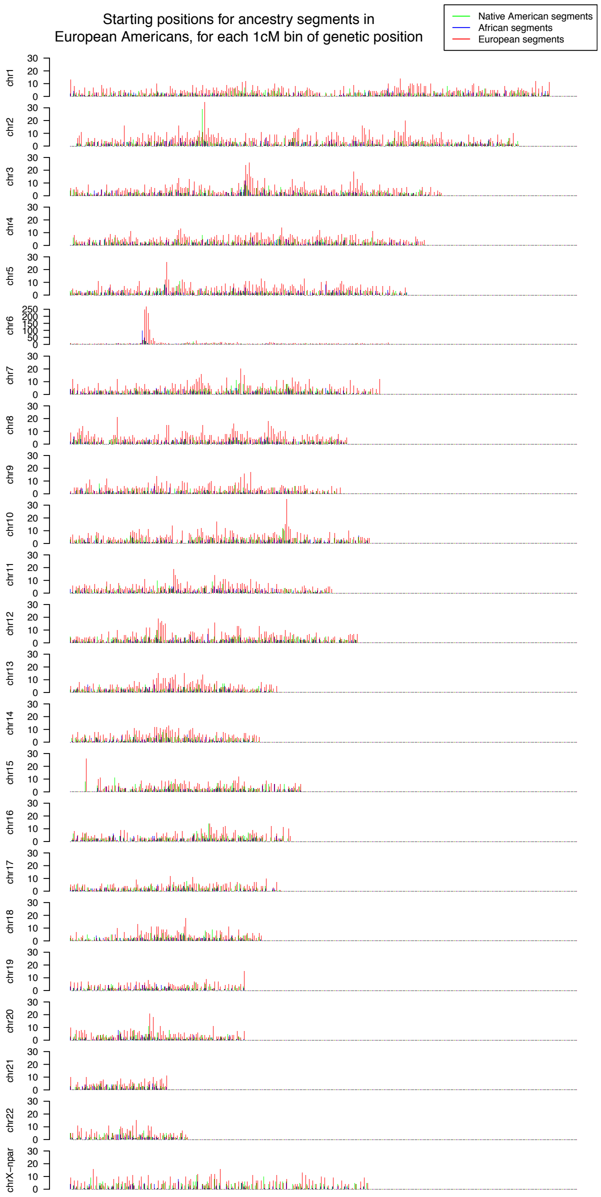


Figure S16: **Distribution of ancestry segment start positions across the genome in self-reported European Americans.** The number of segments that start within a 1cM position along the genome, for each chromosome, are shown by a vertical bar, colored corresponding to African (blue), European (red) or Native American (green) ancestry. Since the vast majority of segments start at the left-most part of each chromsome, the first 5cM of each chromosome are omitted from each plot.

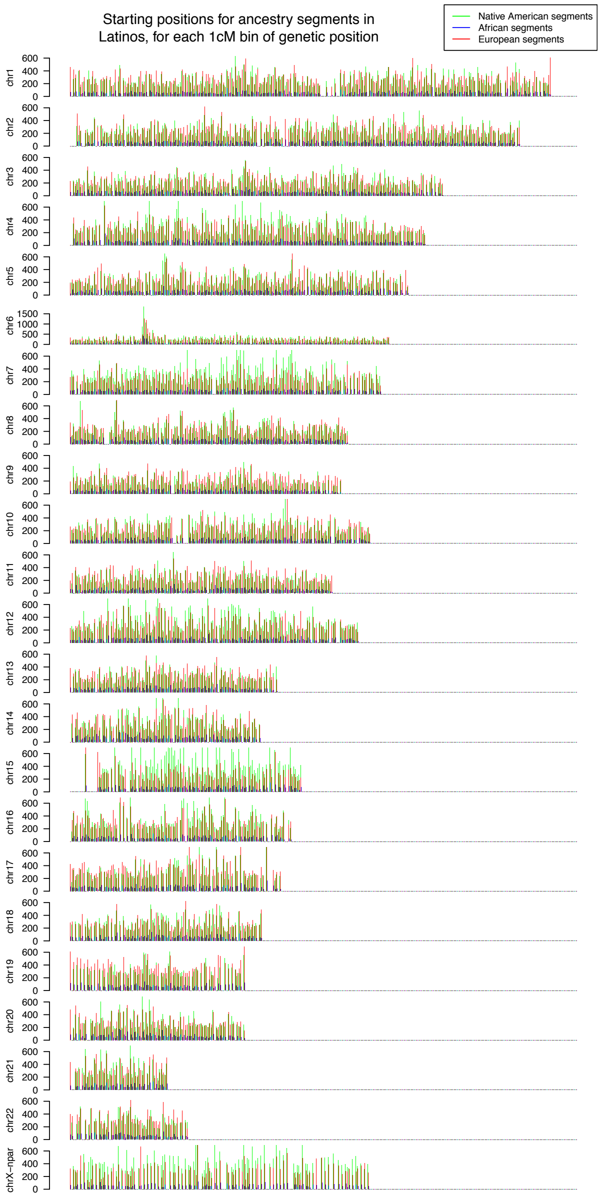


Figure S17: **Distribution of ancestry segment start positions across the genome in self-reported Latinos.**The number of segments that start within a 1cM position along the genome, for each chromosome, are shown by a vertical bar, colored corresponding to African (blue), European (red) or Native American (green) ancestry. Since the vast majority of segments start at the left-most part of each chromsome, the first 5cM of each chromosome are omitted from each plot.

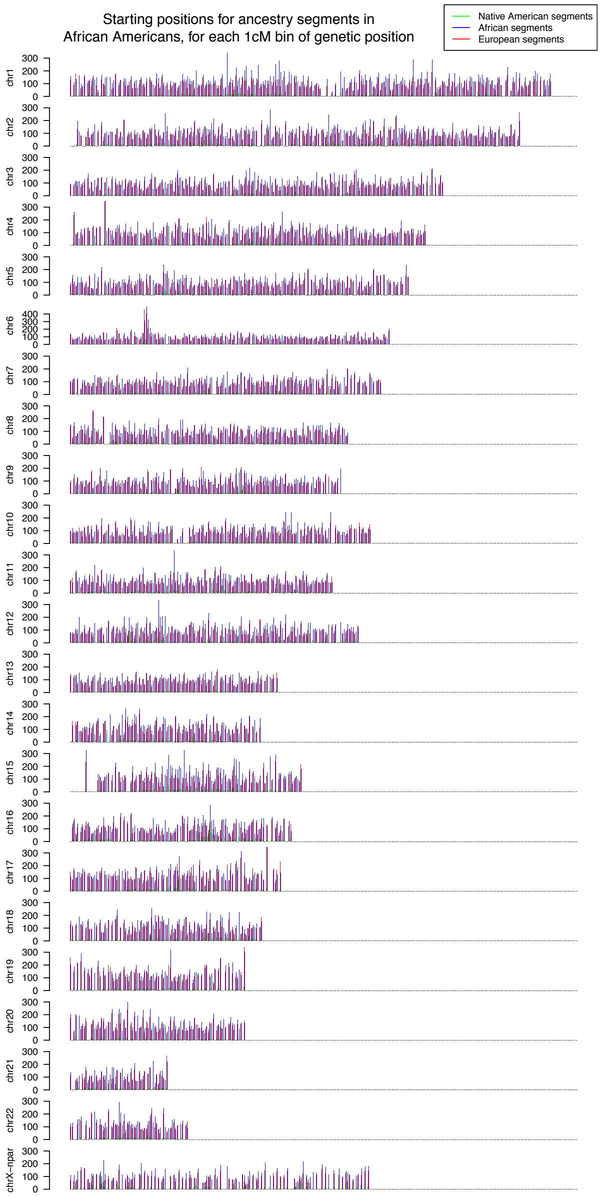


Figure S18: **Distribution of ancestry segment start positions across the genome in self-reported African Americans.**The number of segments that start within a 1cM position along the genome, for each chromosome, are shown by a vertical bar, colored corresponding to African (blue), European (red) or Native American (green) ancestry. Since the vast majority of segments start at the left-most part of each chromsome, the first 5cM of each chromosome are omitted from each plot.
